# REV-ERBβ Binding Pocket Dynamics with Implications for Rational Design of Small Molecule Modulators

**DOI:** 10.1101/2024.04.13.589008

**Authors:** Shriyansh Srivastava, A.M. Vishnu, Rakesh Thakur, Ashutosh Srivastava

## Abstract

REV-ERBβ is a nuclear receptor (NR) with heme as an endogenous ligand that regulates its transcriptional activity. With key role in cellular functions such as glucose metabolism, immune response, and dysregulation in pathologies such as Type-2 diabetes mellitus and obesity, small molecule agonists and antagonists targeting REV-ERBs have been discovered. However, due to lack of crystal structures in complex with these compounds, the structural and dynamical basis of these activities still remains elusive and hinders rational design of molecules targeting REV-ERB. Using molecular dynamics simulations and docking studies, we have characterized the dynamics of REV-ERBβ ligand-binding domain (LBD) in different conformational states. The presence of heme in the binding pocket within LBD was found to stabilize its dynamics as well as nuclear co-repressor (NCoR) peptide binding. We further show that the binding of antagonist destabilizes the NCoR peptide binding to LBD mediated by loss of interactions with residues at the NCoR-REV-ERBβ interface. These findings could be utilized to design molecular scaffolds with better activity and selectivity against REV-ERBβ.

## 1. Introduction

Nuclear receptors (NRs) belong to a class of receptors that mediate their action *via* modulation of transcription of the target genes. REV-ERBα and REV-ERBβ belonging to the nuclear receptor subfamily 1, group D (referred to as NR1D1 and NR1D2) act as transcriptional regulators by binding to the same DNA response element (consensus sequence-AGGTCA) as the Retinoic Acid Receptor (RAR) - Related Orphan Receptors (ROR) (Burris et al., 2023; Weikum et al., 2018). REV-ERBα and REV-ERBβ have been reported to regulate the expression of genes involved in crucial cellular functions such as circadian rhythms, metabolism, and immune system (Kojetin & Burris, 2014). The role of REV-ERBs is central to regulate the expression of other core-clock proteins such as Brain and Muscle ARNT-Like 1 (BMAL1), which is activated by the ROR transcription factor. REV-ERBβ competes with ROR to bind the related orphan receptors element (RORE) binding sites of DNA to inhibit the transcription of BMAL1 (Fig. 1A) (Guillaumond et al., 2005; Ikeda et al., 2019).

**Figure 1:**
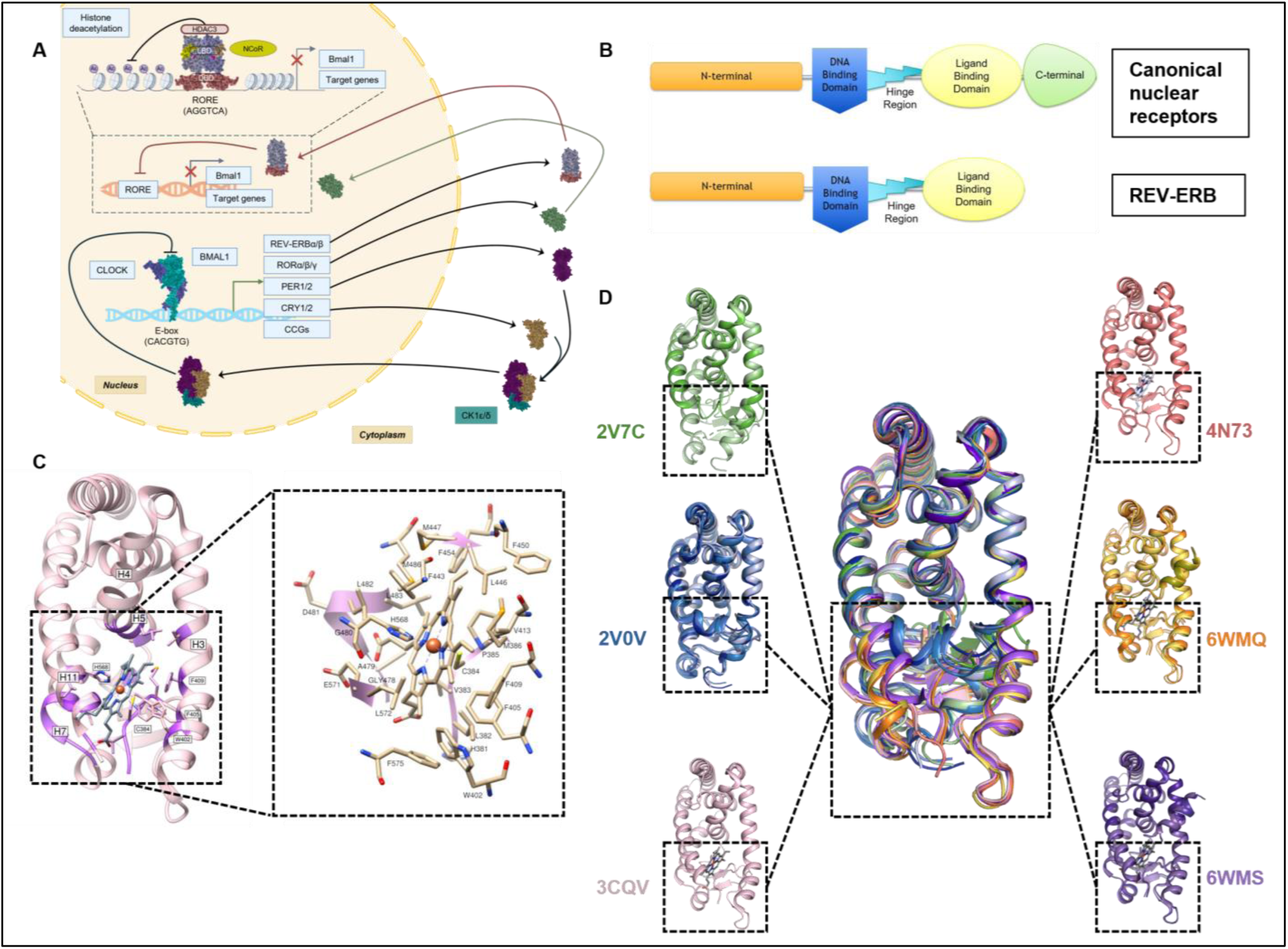
Role of REV-ERBβ as a ligand-dependent transcriptional regulator. **(A)** Schematic illustration of mammalian circadian transcription-translation feedback loop (TTFL) formed by core-clock proteins: Circadian Locomotor Output Cycles Kaput (CLOCK), BMAL-1, Cryptochrome (CRY1/2), Period (PER1/2), ROREα/β/γ, and REV-ERBα/β. Protein structures for representation in the schematic downloaded from PDB; CLOCK-BMAL1 heterodimer (cyan and blue, PDB ID: 4F3L), CRY1 (golden, PDB ID: 4K0R), PER1 (dark magenta, PDB ID: 4DJ2), RORγ (light green, PDB ID: 3KYT), casein kinase (CK) 1δ (dark green, PDB ID: 4JJR), REV-ERBα DNA binding domain (red, PDB ID: 1GA5), and REV-ERBβ ligand-binding domain (light blue, PDB ID: 6WMQ (heme and NCoR ID1 peptide removed)). Regulatory function of REV-ERBα/β is highlighted in the black box with the detailed view of REV-ERBβ functional dimer comprising heme, NCoR, HDAC3, and RORE response element (schematic protein assembly constructed by combining REV-ERBα DNA binding domain (red, PDB ID: 1GA5), and REV-ERBβ ligand-binding domain structures (light blue, PDB ID: 6WMQ)). Protein structures rendered in PyMOL. **(B)** Comparison of LBD architecture of canonical nuclear receptors and REV-ERB. Crystal structure of heme-bound REV-ERBβ LBD (PDB ID: 3CQV) with key binding site residues and helices are highlighted (heme iron-center denoted by orange sphere, coordinated with C384 and H568 located in the N-terminal loop and Helix-11, respectively). **(D)** Conformational diversity in REV-ERBβ LBD highlighted by crystal structures in Apo (PDB ID: 2V7C (green) and 2V0V (blue)), heme (PDB ID: 3CQV (pink)), cobalt porphyrin (PDB ID: 4N73 (red)) and heme+ID peptide (PDB ID: 6WMQ (orange) and 6WMS (purple)) bound states. Conformational differences in the binding pocket region are highlighted with black boxes. Figures **(C)** and **(D)** were rendered in Chimera and PyMOL, respectively.

REV-ERBα and REV-ERBβ follow the domain architecture of NR superfamily of proteins with N-terminal DNA binding domain, followed by a linker and a C-terminal ligand binding domain. The ligand binding domain (LBD) possesses the binding pocket for ligands known to bind REV-ERBα/β (Fig. 1B). The binding pocket comprises residues in the N-terminal loop, Helix-3, Helix-5, β-sheet, Helix-7, and Helix-11 (Fig. 1C) (Weikum et al., 2018). Compared to other members of NR superfamily, REV-ERBs lack Helix-12 in the C-terminal region. Helix-12 is critical for the recruitment of coactivator peptides to promote transcription of target genes (Burris et al., 2023). Thus, REV-ERBs can only function as transcriptional repressors of their target genes. Heme binds to REV-ERBα/β as the endogenous ligand, recruits the nuclear corepressor protein (NCoR) and together, the complex represses the transcription of downstream genes. NCoR recruitment is governed by two interacting domain motifs (ID1 and ID2) present in the NCoR receptor interaction domain (RID) and the complex further recruits histone deacetylases (HDAC-3), resulting in transcriptional repression (Mosure et al., 2021; Raghuram et al., 2007; Weikum et al., 2018).

The role of REV-ERB has been investigated in diverse cellular functions and its dysregulation has been reported in several pathologies, making it an attractive drug target. Efforts have been made to experimentally discover small molecules modulating the REV-ERB activity (Uriz-Huarte et al., 2020). The small molecules discovered have been pharmacologically used to regulate the REV-ERB activity in several disease models such as Alzheimer’s, Duchenne Muscular Dystrophy (DMD), type-2 diabetes mellitus (T2DM), obesity, HIV replication cycle, non-alcoholic fatty liver disease (NAFLD), and cancer (Borrmann et al., 2020; Gomatou et al., 2023; Griffett et al., 2020, 2022; Roby et al., 2019; Solt et al., 2012; Y. Wang et al., 2015; Welch et al., 2017; Yuan et al., 2019). While most studies have reported the role of small molecules on α isoform, only a few studies have looked at the β isoform (De Mei et al., 2015; Torrente et al., 2015; Uriz-Huarte et al., 2020; S. Wang et al., 2020). The ligands have primarily been discovered using fluorescence resonance energy transfer (FRET) and cellular gene reporter assays (Grant et al., 2010; Rasmussen et al., 2022). Though the effect of these compounds has been studied *in vitro* and *in vivo*, their binding modes and molecular basis have not been established due to the lack of experimentally derived complex structures.

The first and the only structure of REV-ERBα in complex with a synthetic ligand STL1267 (PDB ID: 8D8I) indicated the binding site of the ligand to be same as heme, although with a different binding mode (Murray et al., 2022). The synthetic ligand was more localized to the hydrophobic regions of the pocket. The NCoR ID1 formed antiparallel β-sheet with C-terminal residues of Helix-11, which was not observed in the heme bound state (PDB ID: 6WMQ) (Mosure et al., 2021; Murray et al., 2022).

Apart from this, no complex structures with synthetic ligands have been resolved for REV-ERBα or REV-ERBβ. In this study, we have characterized the residue level differences in binding of co-repressor peptides as a consequence of agonist or antagonist binding to REV-ERBβ. We have analyzed the differences in the crystal structures of REV-ERBβ LBD in presence or absence of a ligand, following which, we used molecular dynamics (MD) simulations to understand the dynamics REV-ERBβ LBD in different states. Lastly, we have performed molecular docking and MD studies on small molecule complexes to understand their effect on NCoR binding.

## 2. Methods

### 2.1 Crystal Structure and Small Molecule Data Curation

The crystal structures for REV-ERBβ LBD (Uniprot accession number Q14995) were retrieved from protein data bank (PDB) (Bateman et al., 2023; Berman, 2000). The details of all the structures are summarized in Supplementary Table S1.

The dataset of small molecules experimentally evaluated against either isoform of REV-ERB protein was retrieved from the ChEMBL database (Gaulton et al., 2012). The SMILES files for compounds associated with REV-ERBα (CHEMBL1961783) and REV-ERBβ (CHEMBL1961784) were downloaded. The compounds were sorted by their pharmacological property, based on the reported literature (Supplementary file 1).

### 2.2 Comparison of Crystal Structures

The crystal structures of REV-ERBβ LBD were split into individual chains resulting in twelve monomeric chains. These monomeric chains were superimposed and pairwise RMSD between the chains were calculated using UCSF-Chimera (Pettersen et al., 2004). Hierarchical clustering was performed based on the RMSD matrix.

Distances were computed between W402, F405, and F409 Cα atoms for 2V0V-A and 3CQV to quantify positional difference in crystal structures using PyMOL measurement wizard. Similarly, distances were also calculated for 2V0V-A and 3CQV crystal between Cα atoms of residues W402-M447 (H3-H5 distance) and W402-H568 (H3-H11 distance).

### 2.3 Identification of binding site residues

REV-ERBβ LBD heme bound crystal structure (PDB ID: 3CQV) was used to identify the binding site residues. The residues within the range of 5.0 Å of the heme present in the crystal structure were selected using zone selection tool in UCSF-Chimera (Pettersen et al., 2004) and identified as binding site residues (Supplementary Table S2A).

### 2.4 Molecular Dynamics Simulations

Molecular Dynamics (MD) simulations were performed using Desmond (Schrödinger Release 2022-3: Desmond Molecular Dynamics System, D. E. Shaw Research, New York, NY, 2022. Maestro-Desmond Interoperability Tools, Schrödinger, New York, NY, 2022). The crystal structures were prepared using Protein Preparation Wizard (ProteinPrep) in Schrödinger with default parameters to replace selenomethionine residues with methionine, remove waters, add missing residue atoms, and minimize the structure (Schrödinger Release 2022-3: Protein Preparation Wizard; Epik, Schrödinger, LLC, New York, NY, 2022). The processed structures were used for setting up the system for MD simulations. System builder tool was used to set up the system for simulations with OPLS4 force field (Lu et al., 2021). The system was simulated in rhombic dodecahedron xy-hexagon box with TIP3P water model as explicit solvent. The system was neutralized with Na^+^ and Cl^-^ ions, and further ions were added to make the ionic concentration of 0.15 M. Molecular Dynamics tool in the Desmond module was used to minimize the energy and equilibrate the system with default parameters.

The final MD run was performed using NPT ensemble at 300K and 1.01325 bar pressure with the option to relax the model system before simulation. Different initial velocities were assigned for each simulation replicate, by deselecting the custom random seed option. The details of all the simulations performed in this study are given in Supplementary Table S6.

### 2.5 Analysis of protein trajectories

The simulation trajectories were analyzed using simulation interactions diagram (SID) analysis tool with default settings present in Schrödinger Desmond module (Schrödinger Release 2022-3: Desmond Molecular Dynamics System, D. E. Shaw Research, New York, NY, 2022. Maestro-Desmond Interoperability Tools, Schrödinger, New York, NY, 2022).

Principal Component Analysis (PCA) was performed on the protein backbone using gmx covar and gmx anaeig. Briefly, the replicates for 2V0V-A, Apo-3CQVΔN-ter, and Apo-3CQV trajectories, comprising 10,000 frames each, were concatenated using gmx trjcat. The concatenated trajectory was fitted on the 0^th^ frame of 2V0V-A and gmx covar was used to construct the covariance matrix, calculate eigenvalues, and eigenvectors. The concatenated protein trajectory was further projected to the eigenvectors, and principal components were calculated with gmx anaeig function.

The distances between Cα atoms of protein were measured using the built-in Desmond measurement tool. For replicates, the respective trajectories were first concatenated, and distances were measured between residue pairs (Supplementary Table S3). For 3CQV, 6WMQ-A, and 6WMS-A trajectories, distances were calculated between the heme iron center (Fe^+3^) with S-atom of C384 and imidazole side chain N-atom of H568.

### 2.6 Molecular Docking

All molecular docking studies were performed in Schrödinger with OPLS4 force field (Schrödinger Release 2022-3: Maestro, Schrödinger, LLC, New York, NY, 2022) (Lu et al., 2021).

To establish the validity of our docking protocol, heme bound REV-ERBβ LBD crystal structure (PDB: 3CQV) was used as control (Pardee et al., 2009). Briefly, the protein was prepared with the ProteinPrep protocol. The binding site residues were identified from crystal structure using the protocol mentioned above to generate the docking grid. The co-crystallized ligand (heme) was removed and further processed by Jaguar module in Schrödinger (Schrödinger Release 2022-3: Jaguar, Schrödinger, LLC, New York, NY, 2022). The quantum mechanical optimization for heme was performed using B3LYP density functional theory and 6-31G** basis set. The energy minimized heme was re-docked in XP mode. RMSD of re-docked pose was calculated using co-crystallized ligand as a reference and interactions were compared.

Molecular docking was performed for agonist and antagonist molecules with the Apo-3CQV structure model. In the preprocessing step, the small molecules were prepared using the LigPrep module and the generated conformers were used for docking. The Apo-3CQV structure was prepared using the ProteinPrep protocol. The energy minimized protein structure was further processed by Receptor-Grid Generation tool to generate the docking grid with the centroid defined by the binding site residues. Standard precision (SP) docking was performed with the agonists and antagonists. Extra precision mode of docking (XP) was performed using top conformers of all the molecules using Ligand Docking tool in the Glide module of Schrödinger (Schrödinger Release 2022-3: Glide, Schrödinger, LLC, New York, NY, 2022). Based on the docking score in 3CQV docked complexes, the top three agonists and antagonists were selected and docked in XP mode to Apo-6WMQ-A and Apo-6WMS-A.

### 2.7 Analysis of Docked Complex Simulation Trajectories

The simulation trajectories were analyzed using simulation interactions diagram (SID) analysis tool. Briefly, to evaluate the dynamics of small molecules, ligand ASL was set to ‘small molecule (res.type UNK)’ with protein set as ‘protein (Chain.name A) and NCoR peptide (chain.name E)’. To analyze the dynamics of NCoR peptides, ligand ASL was set to ‘chain.name E’ with protein set as ‘chain.name A’ in the SID analysis tool selection. RMSF for Cα atoms of NCoR peptide (ID1/2) and LBD in different states was calculated to assess the residue-wise fluctuations across the simulation trajectories.

### 2.8 ID1/2 Peptide Binding Affinity Calculations from MD Simulation Trajectories

51 equally spaced frames were extracted from the last 100ns of heme bound (6WMQ-A and 6WMS-A) and last 40ns of agonist/antagonist bound trajectories (Apo-6WMQ-A-agonist, Apo-6WMQ-A-antagonist, Apo-6WMS-A-agonist, and Apo-6WMS-A-antagonist). Binding affinity for ID peptide (chain ‘E’ in complex) with REV-ERBβ LBD (chain ‘A’ in complex) in different states was calculated using PRODIGY (PROtein binDIng enerGY prediction) standalone package (Vangone & Bonvin, 2015). Briefly, chain A and chain E were selected for binding affinity calculations at 27°C. Non-parametric Mann-Whitney-Wilcoxon two-sided test was performed to compare the binding affinity values in different states using statannotations python package (Charlier et al., 2023).

### 2.9 Protein Contact Network Construction and Network Analysis

Protein contact network (PCN) is a graph theoretic representation of 3D structure of a protein where nodes are usually defined by Cα atoms of amino acids and edges represent interatomic distance between Cα atoms based on a custom distance cutoff. The distance matrix is used to generate an adjacency matrix which takes values 0 and 1 based on the distance between the Cα atom pairs. The Cα atom based PCN have been previously constructed at an interatomic distance of ≤7 Å which have successfully captured residue-level interactions to understand crucial information such protein structure-function relationships, impact of mutations, and allosteric communications (Srivastava & Sinha, 2014, 2017).

We therefore constructed an undirected and weighted PCN with frames used in binding affinity analysis at a cutoff of ≤7 Å. The adjacency matrix for each frame was generated where A_ij_ = 1, if Cα atom pair lie within ≤7 Å distance, else A_ij_=0 (if distance >7 Å and i=j, where i and j represent amino acid Cα atoms). The edge weights were calculated by summating the adjacency matrix for 51 frames followed by normalization. PDB files for each frame were parsed using Bio.PDB module from Biopython package (Cock et al., 2009). PCN generation and downstream analysis were performed using in-built functions in python libraries such as NumPy, Pandas, and NetworkX (Hagberg et al., 2008).

We next performed difference Contact Network Analysis (dCNA) developed by Hamelberg and colleagues (Yao et al., 2018). Briefly, dCNA takes two or more conformational ensembles as input and constructs a community-community difference network where each community is a node and the interaction strength between them is the sum of weights of all the edges connecting two communities. We employed this method to calculate a community-community difference network for Apo-6WMQ-A-agonist/antagonist and Apo-6WMS-A-agonist/antagonist with the distance cutoff (dc) set to ≤7 Å, and the contact probability cutoff (Pc) set to 0.7. The rest of the parameters were kept default.

## 3. Results and Discussion

### 3.1 Crystal Structures of REV-ERBβ LBD Highlight Conformational Variability of Binding Pocket

Crystal structures of REV-ERBβ LBD have been resolved in three different states: Apo, heme-bound, and heme+NCoR bound (Fig. 1D and Supplementary Table S1). The endogenous ligand, heme, binds to the LBD of REV-ERBβ by forming coordination bonds with its iron center (Fe^+3^) and residues C384 and H568 on either side of the binding pocket, at the N-terminal loop and Helix-11, respectively. In addition, the binding of heme is stabilized by other interactions at the N-terminal loop (H381, L382, V383, P385, M386), Helix-3 (W402, F405, F409, V413), Helix-5 (F443, L446, M447, F450), β-sheet (F454), loop before Helix-7 (G478, A479), Helix-7 (G480, L482, M486), Helix-11 (E571, L572), and C-terminal loop (F575) (Fig. 1C and Supplementary Table S2A). This suggests that N-terminal loop, Helix 3, Helix 5, β-sheet, Helix 7, Helix 11 and C-terminal loop play an important role in binding of ligand in the LBD (Pardee et al., 2009).

The NCoR binds to a REV-ERBβ dimer through ID1 and ID2 with each monomeric REV-ERBβ LBD subunit, respectively (Mosure et al., 2021). The residue, K421, located on Helix-3, is critical for NCoR recruitment and interaction stabilization, supported by additional non-bonded contacts between REV-ERBβ LBD and ID1/2 peptides predominantly with Helix-3 to 5 (Supplementary Fig. S1A-D and Supplementary Table S2B-C) (Laskowski et al., 2018).

To quantify the conformational differences between the crystal structures of REV-ERBβ LBD, we superimposed the individual chains of all the crystal structures and clustered them based on RMSD. The twelve chains corresponding to six PDB crystal structures formed two distinct clusters for apo and bound form (Fig. 2). The bound cluster further formed distinct sub-clusters based on heme or heme+NCoR bound state (Fig. 2A-B). Considerable differences in Helix-3 and 11 were seen, wherein, helices in apo form were bent inwards to occupy the vacant heme binding pocket. The position of binding site residues W402, F405, and F409 in Helix 3 were distant by 11.5Å, 8.5Å, and 3Å in heme bound form (3CQV) from their position in apo form (2V0V-A) (Fig. 2C). Furthermore, the distance between Cα atoms of W402-M447 (hereafter referred to as H3-H5 distance) for 2V0V-A and 3CQV were 15.3Å and 20.2Å, respectively. The distance between Cα atoms of W402-H568 (hereafter referred to as H3-H11 distance) for 2V0V-A and 3CQV were 13.2Å and 16.6Å (Fig. 2D). These observations combined with the clustering suggested that the binding pocket of REV-ERBβ LBD could take up different conformations based on the binding state.

**Figure 2:**
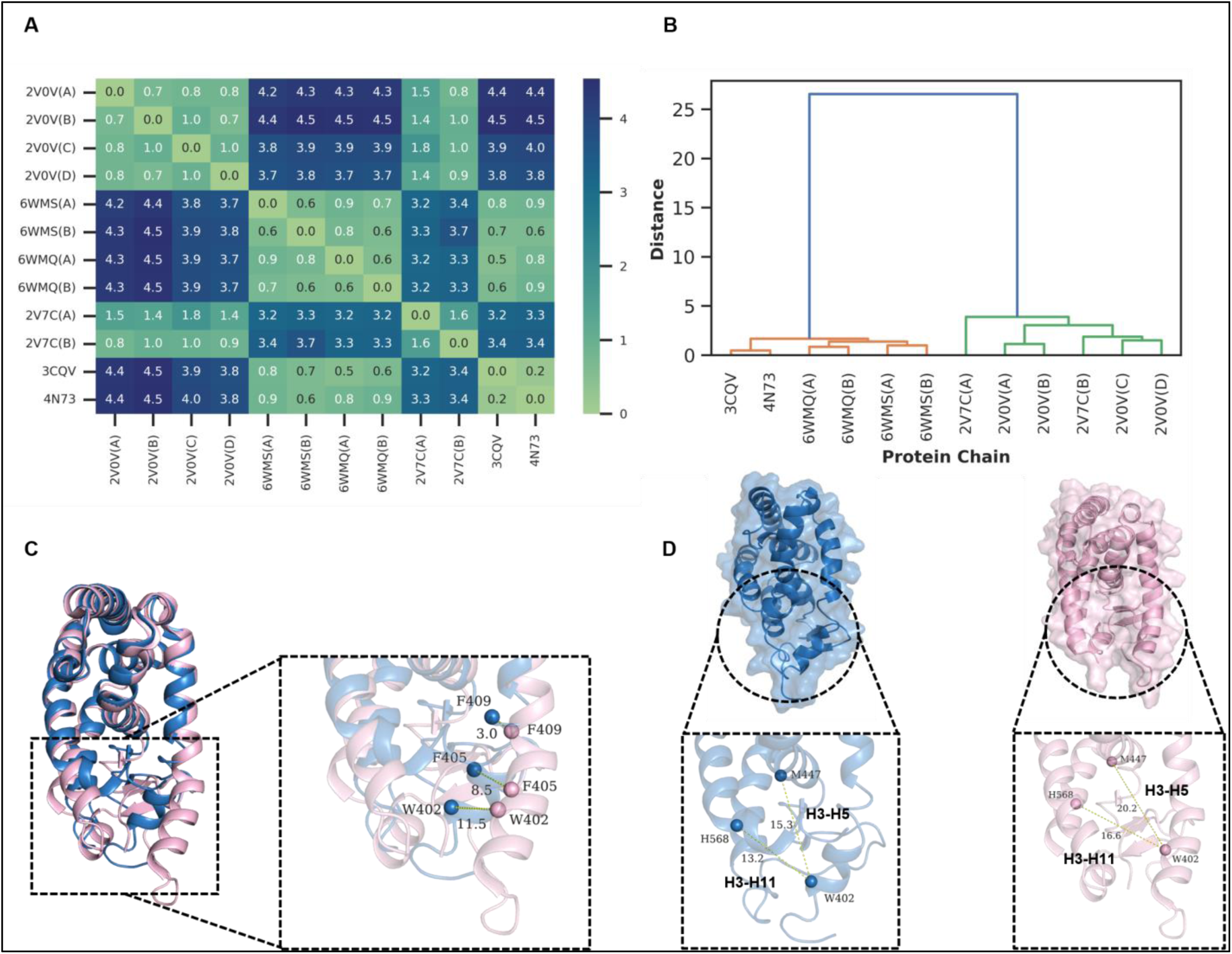
Characterization of REV-ERBβ LBD conformation in different states. **(A)** Heatmap depicting pairwise Cα-atom RMSD values between individual chains of REV-ERBβ LBD resolved in crystal structures **(B)** Hierarchical clustering of REV-ERBβ LBD chains based on pairwise RMSD matrix. Orange cluster represents heme and heme+ID peptide bound REV-ERBβ LBD chains, while chains in Apo-form are denoted by the green cluster. **(C)** Positional differences of binding site residues (W402, F405, F409) in Helix-3 in Apo (PDB ID: 2V0V-A, blue) and heme-bound (PDB ID: 3CQV, pink) states. **(D)** Surface representation of REV-ERBβ LBD in Apo and heme-bound states with detailed view of distances between H3-H5 and H3-H11 in Apo and heme-bound states highlighting conformational differences. Distances were measured in Å. Heme molecule from 3CQV crystal structure removed for clarity. Figures rendered in PyMOL.

### 3.2 Molecular Dynamics Simulations Reveal Conformational Flexibility of Ligand Binding Pocket

To understand dynamics of the binding site in Apo state, we performed three independent 1µs MD simulations of 2V0V-A. The RMSD of Cα atoms for the replicates remained stable around 2-5Å, with some deviations in the beginning and towards the middle of the second replicate (Supplementary Fig. S2A and Supplementary movie 1). Similarly, the radius of gyration for the simulation trajectories were found to be stable between ∼16.5-19Å (Supplementary Fig. S2B). To further investigate the flexibility in different regions of protein, we calculated RMSF for the protein trajectory. Considerable fluctuations were observed for the residues in the loop regions, Helix-3, β-sheets, Helix-11, and C-terminal region including the heme binding site residues in Helix-3 (W402, F405, F409, V413) and Helix-11 (H568, E571, L572) (Fig. 3A).

**Figure 3:**
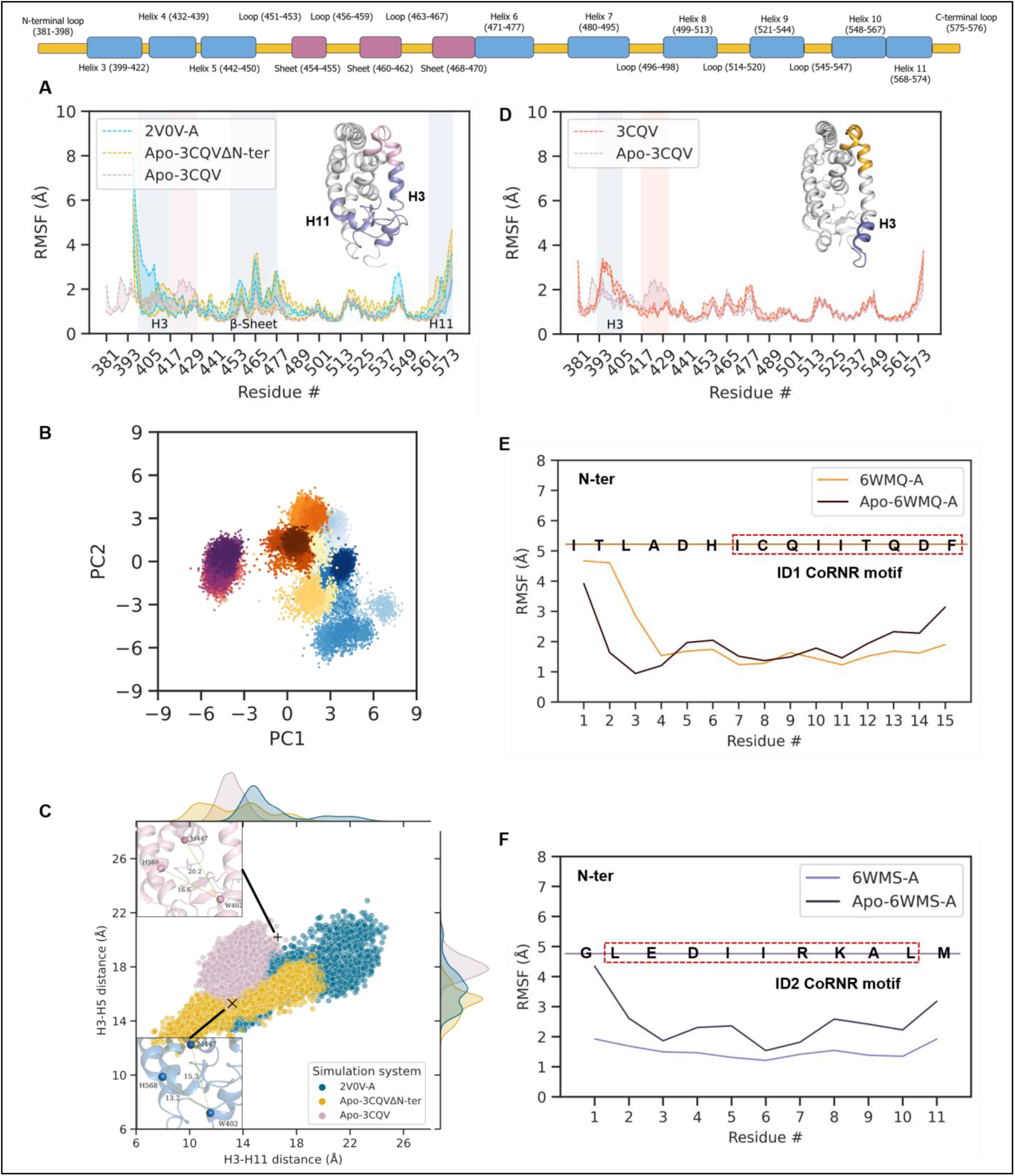
Dynamics of REV-ERBβ LBD and ID peptide in apo and heme bound states. **(A)** RMSF plot for REV-ERBβ LBD Cα atoms highlighting flexibility of the binding pocket regions in absence of heme (2V0V-A: blue, Apo-3CQVΔN-ter: yellow, and Apo-3CQV: pink color). Regions shaded in blue, and pink represent higher fluctuations in 2V0V-A, Apo-3CQVΔN-ter, and Apo-3CQV, respectively. Corresponding regions are colored in the structure. LBD architecture is provided as a 2D schematic for reference. **(B)** Projections of concatenated simulation trajectory (2V0V-A, Apo-3CQVΔN-ter, and Apo-3CQV, see methods) on PC1-PC2 space. The color gradient represents the timeline of trajectories (2V0V-A timeline depicted by transition from light to dark blue, Apo-3CQVΔN-ter timeline depicted by transition from yellow to brown, and Apo-3CQV timeline depicted by transition from red to purple color). **(C)** Distances between Cα atoms of W402-M447 (H3-H5) and W402-H568 (H3-H11) in 2V0V-A, Apo-3CQVΔN-ter, and Apo-3CQV simulation trajectories. Distances in 2V0V-A and 3CQV crystal structure are denoted by (×) and (+), respectively. **(D)** RMSF plot for REV-ERBβ LBD Cα in presence and absence of heme (3CQV: orange, and Apo-3CQV: pink). Regions shaded in blue, and orange represent higher fluctuations in 3CQV and Apo-3CQV, respectively. RMSF plot of **(E)** ID1 and **(F)** ID2 peptide Cα atoms in presence (6WMQ-A and 6WMS-A) and absence (Apo-6WMQ-A and Apo-6WMS-A) of heme. The ID peptide residues are shown in the RMSF plots with CoRNR motif highlighted in red boxes.

Notably, across all the replicates we did not observe any binding pocket conformation similar to the heme bound state (Supplementary Fig. S2C-D). This could be due to the absence of N-terminal loop residues in the 2V0V-A structure (residues: 381-395), a high energy barrier between the apo and bound conformational states, or an induced fit mechanism of heme binding to LBD. To further explore this, we performed MD simulations of the heme bound crystal structure in two different forms. First, after removing heme and N-terminal loop (hereafter referred to as Apo-3CQVΔN-ter) and second, after removing heme only (hereafter referred to as Apo-3CQV). The RMSD and radius of gyration for Apo-3CQVΔN-ter replicates ranged between 2-4.5Å and ∼17-19Å, respectively (Supplementary Fig. S2E-F and Supplementary movie 2a). Similar to 2V0V-A replicates, we did not observe any binding pocket conformations resembling the heme bound state in Apo-3CQVΔN-ter trajectory (Supplementary Fig. S2G-H). The RMSD and radius of gyration values for Apo-3CQV replicates remained stable around 1-3Å and ∼17.5-19Å, respectively, with minor deviations in binding pocket conformation (Supplementary Fig. S2I-L and Supplementary movie 2b). We also observed that Apo-3CQVΔN-ter exhibited similar flexibility to 2V0V-A in most regions of the protein except the β-sheet region, whereas Apo-3CQV showed variations, with a higher flexibility in Helix-3 C-terminal, adjacent loop, and Helix-4 N-terminal regions (residues: 417-432, pink band in Fig. 3A), including the residues critical for recruitment of co-repressor peptide (V417, E418, K421, and Q431) (Fig. 3A and Supplementary Fig. S3A) (Mosure et al., 2021).

We further performed Principal Component Analysis (PCA) on the combined trajectories (see methods). Projection on PC1-PC2 space (with 54.56% of variance explained) revealed that 2V0V-A and Apo-3CQVΔN-ter trajectories occupy similar regions in this space (Fig. 3B, shown in blue and yellow-brown color, respectively; and Supplementary Fig. S3B). In comparison, Apo-3CQV occupied a different space with a smaller spread suggesting lower variations in this trajectory (Fig. 3B, shown in magenta color). The global motions across PC1-PC2 space predominantly comprised of Helix-3 N-terminal, Helix-11 and C-terminal loop regions, including the key binding site residues (W402, F405, H568, E571, L572, and F575) (Supplementary Fig. S3C-E). These observations combined with visual inspection of the trajectories suggested that in the absence of N-terminal loop and heme, the binding pocket takes up conformations similar to apo-form. In addition, it should be noted that presence of N-terminal loop is critical for heme binding, since it harbors C384 residue, essential for coordinate bond formation with heme iron center (Pardee et al., 2009).

To further quantify the conformational variability in the binding pocket, we calculated the distances between the Cα atom of binding site residue pairs on either side of the pocket across all replicates (see methods). The residue pairs used for distance calculations were located in Helix-3, Helix-5, Helix-7, and Helix-11 (W402, F405, L446, M447, F450, G480, L483, M486, H568, and L572) (Supplementary Table S3). Notably, we observed higher variations in the distances in 2V0V-A and Apo-3CQVΔN-ter trajectories as compared to Apo-3CQV trajectories (Fig. 3C and Supplementary Fig. S3F). For example, the H3-H11 distances for 2V0V-A and Apo-3CQVΔN-ter ranged between ∼12-24Å and ∼7-20Å, respectively. However, these distances only ranged between ∼10-17Å in Apo-3CQV trajectory (Fig. 3C).

The above results indicate that the REV-ERBβ LBD exhibits considerable conformational flexibility in the absence of heme and the N-terminal region.

### 3.3 Heme along with N-terminal Residues of LBD Stabilize Binding Pocket

REV-ERBβ LBD MD simulations in Apo, Apo-3CQVΔN-ter, and Apo-3CQV forms revealed the dynamics of ligand binding pocket in different states. In absence of the endogenous ligand and the N-terminal loop, the conformationally open pocket transitioned into a closed apo state, highlighting the heme-induced conformational changes. Interestingly, only removing the heme from the binding pocket did not lead to complete closure of the pocket. This could be due to the N-terminal loop stabilizing the binding pocket. However, considering N-terminal loop is required for binding of heme and is not present in any apo structure, its presence in the binding pocket may be dependent on heme. To further understand this, we performed MD simulation for 3CQV crystal structure (with heme bound).

The heme iron center remained coordinated throughout the trajectory with C384 and H568 and the side chain interactions were stabilized by binding site residues (Fig. S3G-H and Supplementary movie 3). To understand the influence of heme on the binding pocket dynamics, we calculated the RMSF and compared it with Apo-3CQV trajectory. Importantly, we found that binding of heme stabilized the Helix-3 C-terminal, adjacent loop, and Helix-4 N-terminal regions (residues: 417-432, orange band in Fig. 3D), including the residues critical for recruitment of co-repressor peptide (V417, E418, K421, and Q431). Additionally, we observed a higher flexibility in the N-terminal loop and Helix-3 N-terminal region (residues: 392-405) (Fig. 3D and Supplementary Fig. S3I). Other than this, we did not observe any major differences between Apo-3CQV and 3CQV trajectories. These results suggest that presence of heme stabilizes the binding pocket dynamics at Helix-3 C-terminal and adjacent regions to govern an overall conformational stability of REV-ERBβ LBD, that may be critical for NCoR peptide recruitment.

### 3.4 Heme confers stability to NCoR peptide interactions with LBD

Nuclear receptors are ligand-dependent transcriptional regulators where ligand binding causes recruitment of co-activator/repressor peptides. Mosure and colleagues reported that the binding of NCoR in presence of heme was governed by interactions of ID1/ID2 peptide with LBD (Fig. S1A-D and Supplementary Table S2B-C) (Mosure et al., 2021). The conserved I/LxxI/LI/LxxxI/L/F co-repressor NR motif (hereafter referred to as CoRNR) interacts with the AF2 surface formed by Helix 3 to 5 (Mosure et al., 2021) (ID1 CoRNR motif: ICQIITQDF and ID2 CoRNR motif: LEDIIRKAL). Previous studies have also reported that residues in Helix-3, 4, 5, and 11 (W402, F409, V413, K414, V417, K421, R427, V435, K439, L572, F575, K576) regulate NCoR binding (Burke et al., 1998; Renaud et al., 2000; Woo et al., 2007).

Based on these observations and our simulations of Apo-3CQV where flexibility in AF2 region increased as compared to 3CQV, we hypothesized that heme binding could stabilize NCoR interactions with the AF2 surface *via* ID1/ID2 peptide. To test this, we performed MD simulations of REV-ERBβ LBD in complex with NCoR ID1 and ID2 peptide with and without heme bound to the binding pocket (PDB ID: 6WMQ, 6WMS, and corresponding apo forms).

Similar to the 3CQV trajectory, heme remained coordinated and bound in the pocket for 6WMQ-A and 6WMS-A trajectories (Supplementary Fig. S4A-B and Supplementary movies 4-5). Interestingly, we observed a higher RMSF for Cα atoms of ID1 peptide residues interacting at the AF2 surface (ITQDF) in Apo-6WMQ-A trajectory compared to its heme-bound counterpart (6WMQ-A) (Fig. 3E). These residues represented a part of CoRNR motif, highlighting increased fluctuations in absence of heme. Similarly, RMSF calculations for ID2 peptide Cα atoms in Apo-6WMS-A and heme-bound counterpart (6WMS-A) trajectories revealed higher fluctuations when heme was removed (Fig. 3F). We further looked at the dynamics of ID1 and ID2 peptides and observed key differences in LBD-ID peptide interactions upon comparative analysis of the trajectories (Supplementary movies 4-7). In the absence of heme (Apo-6WMQ-A), we found loss of ID1 peptide interactions with key AF2 surface residues (K421, F426) and slight decrease in interactions with residues at Helix-3 and 4 (K414, V417, L438) than in heme-bound trajectory (6WMQ-A). Further, the interactions of ID1 peptide N-terminus with Helix-10, Helix-11 and C-terminal were prominent throughout the Apo-6WMQ-A trajectory but were lost in the later part of heme-bound trajectory (6WMQ-A) (Supplementary Fig. S4C-D). Previous studies have reported differences in the binding mode of ID1 N-terminus, where it formed antiparallel β-sheets in Apo REV-ERBα isoform LBD crystal structure (PDB ID: 3N00) (Phelan et al., 2010). This sheet formation was lost in structures of REV-ERBβ LBD co-bound with heme and ID1 peptide, highlighting difference in ID1 N-terminus interactions upon ligand binding (Mosure et al., 2021). The interactions of ID2 peptide remained similar at AF2 surface in both trajectories (Apo-6WMS-A and 6WMS-A) (Supplementary Fig. S4E-F). However, a slight increase in interactions with residues V417, K421 and decrease with residues F409, K414 were observed in absence of heme. Overall, the dynamics of ID peptides in presence or absence of heme proved the role of heme in stabilizing the LBD-NCoR complex interactions at the AF2 surface and revealed the dynamic behavior of ID1 N-terminus.

### 3.5 Differential effect of Agonists and Antagonists on interaction between NCoR and REV-ERBβ LBD

Both REV-ERB α and β have emerged as potential drug targets in diverse diseases (Kojetin & Burris, 2014). Interestingly, both agonists and antagonists have been discovered for REV-ERB (Uriz-Huarte et al., 2020). As per the current understanding both agonists and antagonists bind REV-ERB LBD at the same (heme) binding pocket. However, the molecular basis of differential mechanisms of the two class of ligands has not been understood. Based on the above results we can hypothesize that the agonists, mimicking the action of heme, would positively affect the NCoR recruitment whereas antagonists would make the NCoR binding weaker.

To investigate this, we performed molecular docking and simulations. The heme bound crystal structure of REV-ERBβ LBD was used to validate our docking protocol (Supplementary Fig. S5A). Experimentally discovered small molecule agonists and antagonists against REV-ERB reported in ChEMBL database were docked to the Apo-3CQV structure (Supplementary file 1). The docking score (hereafter referred to as Dscore) ranged from –10.57 to 0.10 for agonists and – 9.76 to –2.29 for antagonists (Supplementary file 1). The top 3 agonists, CHEMBL2030069, CHEMBL2059573, and CHEMBL2030068 (hereafter referred to as Agonist 1, 2 and 3, respectively) had comparable docking scores but exhibited different interactions (Supplementary Fig. S5B and Supplementary file 1). A similar differential interaction trend was seen for the top 3 antagonists: CHEMBL3590561, CHEMBL3590571, and CHEMBL3590573 (hereafter referred to as Antagonist 1, 2 and 3, respectively) (Supplementary Fig. S5C and Supplementary file 1). Molecular dynamics simulations of these docked complexes revealed that the ligands were flexible but remained bound within the pocket (Supplementary Fig. S6 and Supplementary movies 8-13). To understand if the binding of agonists and antagonists differentially affects the NCoR binding, we first docked the top three agonists and antagonists to Apo-6WMQ-A and Apo-6WMS-A (Supplementary file 1). Based on the Dscore, we focused on Agonist 2 and Antagonist 1 docked complexes (hereafter referred to as agonist and antagonist, respectively) (Supplementary Fig. S7). It is noteworthy that the agonist and antagonist were structurally different, belonging to the class of tertiary amine and diarylalkylamine derivatives, respectively (Supplementary Fig. S7A-B) (De Mei et al., 2015; Shin et al., 2012; Torrente et al., 2015). Interestingly, the antagonist identified through molecular docking belongs to the experimentally discovered series of dual inhibitors targeting REV-ERBβ and autophagy (De Mei et al., 2015; Torrente et al., 2015). We next performed MD simulations for Apo-6WMQ-A and Apo-6WMS-A complexed with agonist or antagonist (here after referred as Apo-6WMQ-A-agonist, Apo-6WMQ-A-antagonist, Apo-6WMS-A-agonist, and Apo-6WMS-A-antagonist). In all the simulations, the small molecules remained within the binding pocket but were flexible (Supplementary Fig. S8). In Apo-6WMQ-A-agonist trajectory, agonist was central to the binding pocket, with dominant interactions in N-terminal loop, Helix-3, 5, 7, 11, C-terminal regions, and ID1 peptide; while antagonist predominantly interacted with residues in Helix-5, 7, 11, and lost interactions with residues in Helix-3 in later part of the Apo-6WMQ-A-antagonist trajectory (Supplementary Fig. S8A-B and Supplementary movie 14-15). In case of Apo-6WMS-A-agonist/antagonist trajectories, agonist interacted with residues in N-terminal loop, Helix-3, 5, 7, 11, C-terminal regions, and ID2 peptide; while antagonist mimicked heme-like conformation and interacted predominantly with residues in N-terminal loop, Helix-3, and Helix-11 (Supplementary Fig. S8C-D and Supplementary movie 16-17).

To understand residue-wise fluctuations in the LBD, we calculated RMSF for the four systems (Apo-6WMQ-A-agonist, Apo-6WMQ-A-antagonist, Apo-6WMS-A-agonist, and Apo-6WMS-A-antagonist) and compared them with their respective Apo and heme-bound trajectories.

We observed higher fluctuations for Cα atoms of LBD in N-terminal loop, Helix-3 N-terminal (residues: 395-410), β-sheets, loop before Helix-6, Helix-6 (residues: 451-480), Helix-11, and C-terminal regions (residues: 568-577) in Apo-6WMQ-A-antagonist trajectory as compared to other systems (Apo-6WMQ-A, 6WMQ-A, Apo-6WMQ-A-agonist) (Fig. 4A). However, in the case of 6WMS trajectories (Apo-6WMS-A, 6WMS-A, Apo-6WMS-A-agonist, Apo-6WMS-A-antagonist), we did not see such differences. A decrease in fluctuations in the N-terminal loop (residues: 381-395) and a minor increase in fluctuations in Helix-11 and C-terminal regions (residues: 568-577) was observed in Apo-6WMS-A-agonist/antagonist trajectories compared to other systems. In addition, Apo-6WMS-A-agonist exhibited higher fluctuations in Helix-3 C-terminal and adjacent loop region (residues: 409-423) (Fig. 4B).

**Figure 4:**
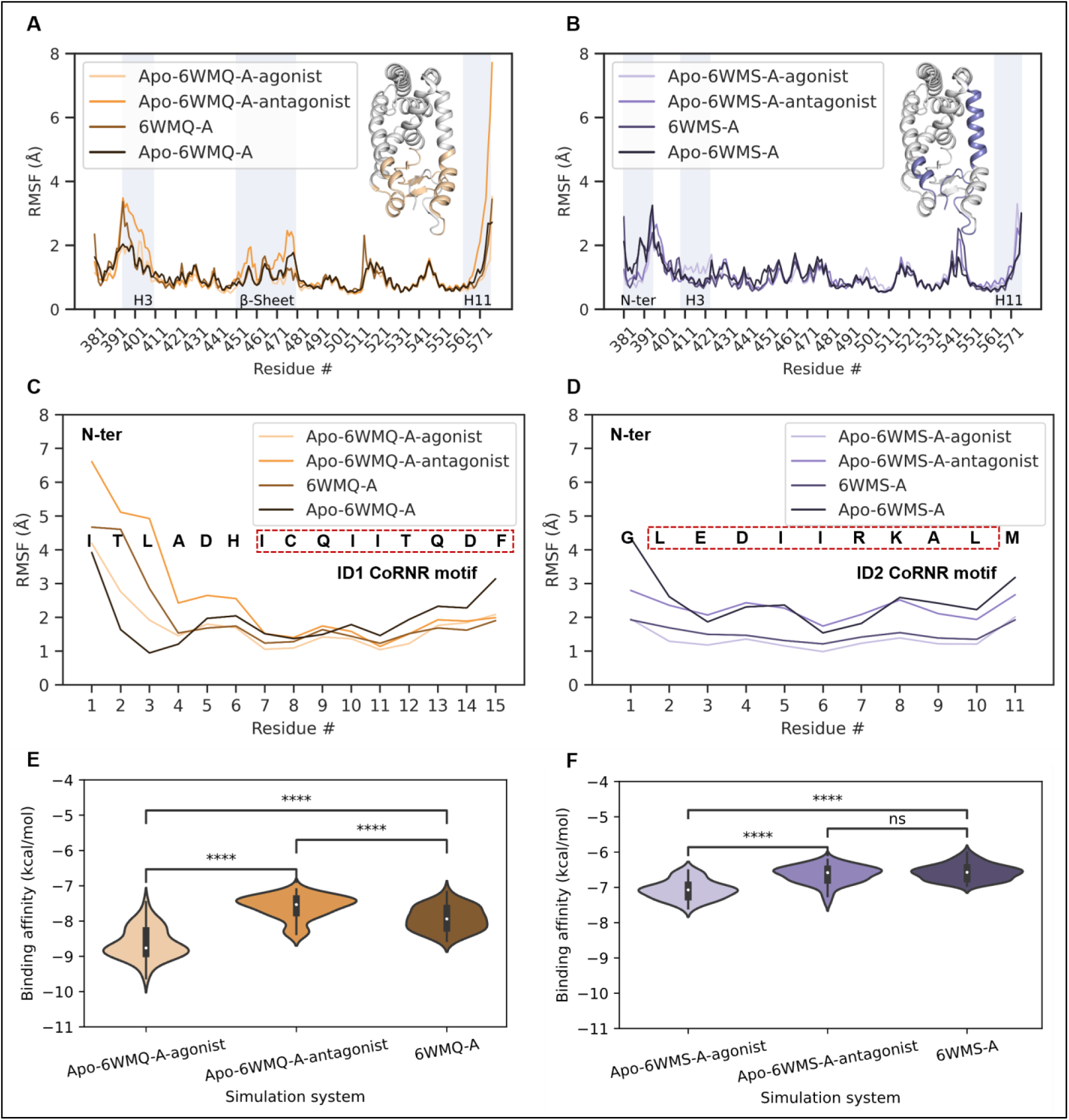
Analysis of Apo-6WMQ-A and Apo-6WMS-A simulation trajectories in presence of agonist and antagonist. Combined RMSF plot for REV-ERBβ LBD Cα atoms from **(A)** ID1 and **(B)** ID2 bound simulation trajectories in presence of agonist, antagonist, heme, and absence of any ligand. Corresponding regions are colored in the structure. RMSF plot of **(C)** ID1 and **(D)** ID2 peptide Cα atoms in presence of agonist, antagonist, heme, and in absence of any ligand. Comparison of binding affinity of **(E)** ID1 and **(F)** ID2 peptide in different simulation systems. The Mann-Whitney-Wilcoxon two-sided statistical test was performed for comparison. **** indicates p-val less than or equal to 0.0001, ns indicates not significant.

We further calculated RMSF of Cα atoms of ID1/2 peptide for all models and compared them with respective Apo and heme bound forms. Importantly, we observed a higher RMSF for ID1 N-terminus (residues: 1-6) (∼2-6 Å) in antagonist bound state, as compared to the counterparts (Fig. 4C). Similarly, RMSF calculations identified a higher fluctuation of ID2 peptide in Apo-6WMS-A-antagonist (Fig. 4D).

From protein-ligand contact map and contact timeline analysis for ID peptides, we could observe loss in interactions of ID1 peptide with residues located in Helix-10, Helix-11, and C-terminal region after ∼50 ns in Apo-6WMQ-A-antagonist compared to Apo-6WMQ-A-agonist trajectory (Supplementary Fig. S8E-F). We did not find such differences upon comparative analysis of Apo-6WMS-A-agonist/antagonist trajectories (Supplementary Fig. S8G-H). Further, it was visible from Apo-6WMQ-A-antagonist trajectory that Helix-11 moved farther as compared to the initial structure (Supplementary movie 15). This visual observation was confirmed by calculating the H3-H11 distance for Apo-6WMQ-A-antagonist trajectory. After ∼50 ns, the distance increased to ∼21 Å and remained in that range. However, such large deviations were not observed in distances for Apo-6WMQ-A-agonist, Apo-6WMS-A-agonist, or Apo-6WMS-A-antagonist trajectories (Supplementary Fig. S7I). These observations suggested that small molecule binding could affect LBD-ID peptide interactions.

To further quantify the effect of structural changes on binding of ID peptide, we calculated ID1/2 peptide binding affinity in respective simulation trajectories, with heme bound trajectory as positive control (see Methods). We observed a significant decrease in ID1/2 peptide binding affinity in presence of antagonist as compared to agonist bound counterparts. The ID1 peptide binding affinity in Apo-6WMQ-A-agonist trajectory ranged –9.62 to –8.76 kcal/mol while it decreased in Apo-6WMQ-A-antagonist trajectory (–8.40 to –7.53 kcal/mol). We observed a comparable ID1 binding affinity value in heme bound 6WMQ-A trajectory (–8.56 to –7.94 kcal/mol) (Fig. 4E). Similarly, ID2 peptide binding affinity in Apo-6WMS-A-agonist ranged – 7.60 to –7.072 kcal/mol which decreased in Apo-6WMS-A-antagonist trajectory (–7.43 to –6.58 kcal/mol). The ID2 peptide binding affinity in heme bound 6WMS-A trajectory ranged –6.95 to – 6.57 kcal/mol (Fig. 4F).

Overall, these combined results suggested that structural changes induced by agonist favored, while the changes upon antagonist binding reduced REV-ERBβ LBD-ID peptide binding affinity.

### 3.6 Network Analysis Reveals Differential ID Peptide Interactions in Agonist/Antagonist Bound Trajectories

To understand the effect of agonist and antagonist binding on interactions between ID peptide and REV-ERBβ LBD at residue-level, we constructed protein contact network (PCN) for trajectory frames in agonist/antagonist bound states (Apo-6WMQ-A-agonist, Apo-6WMQ-A-antagonist, Apo-6WMS-A-agonist, and Apo-6WMS-A-antagonist). The presence of an edge in the network represents that the two residues are closer and hence potentially interacting with each other. We focused our analysis on the edges present at the interface between ID peptides and the REV-ERBβ LBD.

The number and frequency of edges present exclusively between ID1/2 peptide and agonist bound REV-ERBβ LBD was higher than antagonist bound REV-ERBβ LBD (Fig. 5A and Supplementary Table S4). Similarly, for the common edges (present both in Apo-6WMQ-A-agonist and Apo-6WMQ-A-antagonist), the frequency decreased significantly in case of Apo-6WMQ-A-antagonist, suggesting weaker interaction between ID peptide and LBD (Fig. 5B and Supplementary Table S5A).

**Figure 5:**
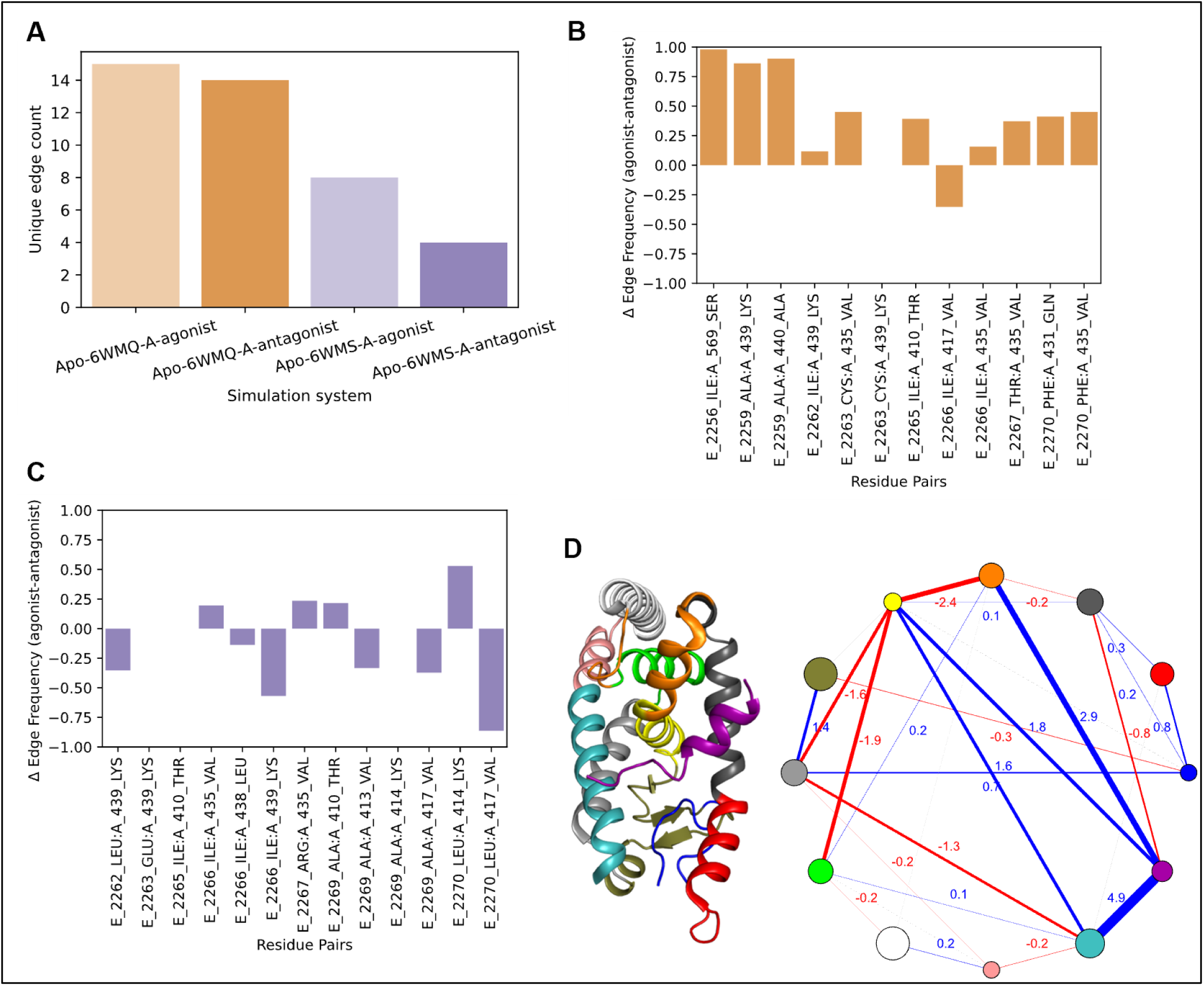
Analysis of Apo-6WMQ-A and Apo-6WMS-A agonist/antagonist protein contact network. **(A)** Comparison of number of unique edges between LBD (chain A) and ID1/2 (chain E) in the presence of agonist/antagonist **(B)** Difference in the edge frequency of the common edges between Apo-6WMQ-A-agonist and Apo-6WMQ-A-antagonist. **(C)** Difference in the edge frequency of the common edges between Apo-6WMS-A-agonist and Apo-6WMS-A-antagonist. Community–community difference contact network of Apo-6WMQ-A-agonist/antagonist (ID1 peptide shown in purple color). Red lines indicate a negative difference, highlighting stronger interactions in the antagonist-bound state, whereas blue lines indicate a positive difference, highlighting stronger interactions in the agonist-bound state. The details of the community membership can be found in Supplementary file 2.

Interestingly, the common edges involved residues from CoRNR motif (I2262-F2270) and AF2 surface (Helix-3 to 5) interactions, and a decrease in the edge frequency further proved the effect of antagonist binding on disrupting the interactions.

Similarly, the Apo-6WMS-A-agonist and Apo-6WMS-A-antagonist frames exhibited 8 and 4 unique edges between ID2 and REV-ERBβ LBD (Fig. 5A and Supplementary Table S4). The unique edges comprised interactions between CoRNR motif and AF2 residues and generally exhibited a lower frequency in both cases. The common edges between Apo-6WMS-A-agonist and Apo-6WMS-A-antagonist frames were also present at the AF2 interacting regions. However, the decrease in edge frequency for antagonist frames were not consistent as observed in ID1 counterpart (Apo-6WMQ-agonist/antagonist) (Fig. 5C and Supplementary Table S5B). These initial results from protein contact network highlighted the direct impact of small molecule binding on ID peptide and REV-ERBβ LBD interactions.

We further investigated if there were global changes in connectivity between the two states by analyzing the community structure in the networks. Community structure analysis has been used widely to understand the protein architecture and its direct relevance on functionality (Delmotte et al., 2011; Srivastava et al., 2020; Srivastava & Sinha, 2014). Difference contact network analysis (dCNA) revealed 12 communities in the Apo-6WMQ-A agonist/antagonist and 10 communities in the Apo-6WMS-A-agonist/antagonist community-community difference contact network pair (Fig. 5D, Supplementary Fig. S9, and Supplementary file 2). In Apo-6WMQ-A, the ID1 peptide formed a separate community (purple) with stronger connections to communities in REV-ERBβ LBD containing residues from Helix 4 (H432-K439; orange), Helix 5 (T442-F450; yellow), Helix 10 and 11 (K557-A574; teal) in agonist bound form as compared to antagonist bound state (Fig. 5D, shown in blue lines). In the case of Apo-6WMS-A, though ID2 peptide formed a separate community, there was no considerable change in its interaction with other communities in the two states (Supplementary Fig. S9). This differential behavior of ID1 and ID2 peptide is in line with our own results as well as previous experimental findings.

With these results, it was evident that agonist binding increased the communication between ID1 peptide and LBD, suggesting that agonist induced favorable structural changes that stabilized the interactions between ID1 peptide and LBD, whereas antagonist binding induced structural changes that disrupted such interactions.

Leveraging the potential of REV-ERB as a drug target has led to the development of small molecules that can regulate its downstream transcriptional activity. While experimental studies have identified modulators of REV-ERB, the structural mechanisms of their pharmacological activity have remained elusive. Using computational methods to understand the structural mechanisms of small molecule activity, we have characterized the residue level changes in LBD and ID peptide interactions upon small molecule binding.

We have first reported the dynamic behavior of REV-ERBβ LBD in the presence or absence of the endogenous ligand, heme. Following the initial observations, we have characterized the role of heme in regulating the dynamics of LBD and the binding of NCoR ID1/2 peptides and lastly, we have reported the differential dynamics of NCoR ID1/2 peptide binding to REV-ERBβ LBD in presence of an agonist or antagonist. The first observation from the MD simulations of apo-forms (2V0V-A, Apo-3CQVΔN-ter, and Apo-3CQV) and heme-bound state (3CQV) highlighted the critical role of N-terminal loop and the endogenous ligand, heme. The binding pocket in Apo-3CQVΔN-ter closed during the simulation and adopted conformations similar to apo-form (2V0V-A), however, presence of N-terminal loop prevented the pocket from closing and increased the pocket stability. It is important to note that the presence of N-terminal loop is critical for heme binding, along with H568 in Helix-11, since it harbors residue C384 essential for heme coordination (Pardee et al., 2009). Of note, heme-bound 3CQV trajectories revealed that presence of both N-terminal and heme could stabilize the region for NCoR recruitment, suggesting that presence of a ligand is essential for the LBD domain to remain in a functional state which is in line with the observed mechanism for other NR ligands (Weikum et al., 2018). Crystal structures (PDB ID: 6WMQ and 6WMS) and experiments have demonstrated the role of endogenous ligand heme to thermodynamically regulate binding of NCoR *via* ID1/2 peptide. The holo-conformation of the LBD was present throughout the 3CQV trajectory, suggesting that ligand binding may follow an induced fit model of binding. In addition, MD simulations of ID peptide bound structures in presence and absence of heme further revealed the dynamic behavior of ID-LBD interactions, and the role of heme to stabilize key interactions for ID1 and ID2 peptides. Understanding these observations highlighted that utilizing the bound form conformation could be more useful for the structure based rational designing of small molecules to target REV-ERBβ. This hypothesis is further supported by our findings from molecular docking and MD simulations of small molecule ligand complexes.

The binding mode of top-ranked agonist (CHEMBL2059573) in our study was in line with a previous computational study investigating binding mode of agonist with REV-ERBα where interactions with Helix-3 phenylalanine residues were favorable for activity (Westermaier et al., 2017). However, the binding mode of antagonist (CHEMBL3590561) in our study exhibited some differences from a recently published study predicting binding of prototypical synthetic antagonist SR8278 with REV-ERBα through GaMD simulations (Rahman & Hegazy, 2024). In our study, the binding mode of top-ranked antagonist was distal to Helix-3, however, GaMD studies revealed significant interactions with Helix-3, Helix-5 and Helix-6 residues in REV-ERBα. The difference in binding mode can be attributed to the difference in the chemical scaffold of antagonists under investigation and the absence of N-terminal loop in REV-ERBα crystal structure (PDB ID: 8D8I) used in the study (Rahman & Hegazy, 2024). Through MD simulations of top-ranked agonist and antagonist from molecular docking screen, we were able to see a direct effect of differences in the binding mode of small molecules on ID1/2 binding dynamics with the LBD. The disruptions of ID1 N-terminus-LBD interactions, an overall higher RMSF for ID2 peptide in presence of antagonist, and differential dynamics induced by agonist and antagonist, evinced the plausible mechanism for the pharmacological activity. In addition, we were able to delineate the residue-level changes in presence of agonist/antagonist by network analysis.

Given that we have biochemical screens for development of newer small molecules, this structural evidence for designing better molecules could be pivotal. This structural understanding is needed because studies have reported contrasting results for a small molecule, that has been identified an agonist in one study, but antagonist in another (Trump et al., 2013; Zhang et al., 2018). Similarly, the activity of SR9009, a prototypical agonist for REV-ERB has recently been reported to cause downstream effects in a REV-ERB independent manner (Dierickx et al., 2019). In addition, with the discovery of newer chemical scaffolds, understanding the structural basis of activity could be an effective tool for optimizing lead compound synthesis.

Lastly, the structural mechanisms identified from the current study can be critical to develop isoform selective REV-ERB modulators that has recently gained interest (He et al., 2023). Even with binding site residues being conserved in the isoforms, the LBD only share ∼71% similarity, which may give rise to functional conformational differences while recruiting NCoR *via* ID1/2 (Mosure et al., 2021; Murray et al., 2022). The conformation of N-terminal loop remains stable in the Apo-3CQV trajectory suggesting that it may have additional role in stabilizing the pocket. From the results of molecular docking and MD simulations of small molecules, we could observe that residues within the loop also interact with small molecules, supporting their functional relevance. Although this study is aimed at investigation of REV-ERBβ LBD, a qualitative comparison with REV-ERBα LBD co-crystallized with STL1267 (PDB ID: 8D8I) suggests that the conformation of N-terminal loop also has direct implications on NCoR binding. The absence of N-terminal loop allows STL1267 to be buried deep into the binding pocket; because of which the ID1 peptide forms an antiparallel β-sheet similar to REV-ERBα Apo form (PDB ID: 3N00) (Phelan et al., 2010). Evaluating these hypotheses would aid in better understanding of the structural mechanisms underlying pharmacological activity. With the crystal structures resolved for both isoforms, a comparative structural study on experimentally discovered small molecules could strengthen the niche to develop isoform selective modulators.

## 4. Conclusion

Previously discovered small molecules modulating REV-ERB activity have been identified using FRET-based and cellular gene reporter assays that could indirectly highlight the regulation of REV-ERB activity with only a few computational studies on REV-ERBα (Rahman & Hegazy, 2024; Westermaier et al., 2017). Our study complements these experimental observations and establishes an understanding on the structural basis of the molecular effect exhibited by small molecule agonists and antagonists. Our findings from molecular docking and MD simulations revealed a difference in the binding modes for agonist and antagonist molecules within the orthosteric binding pocket that could lead to differential effect on the interactions of NCoR ID peptides with REV-ERBβ LBD.

## ASSOCIATED CONTENT

### Supporting information description

Supplementary Material File: Supplementary figures and tables associated with the study

Supplementary file 1:SMILES and details of molecular docking results

Supplementary file 2: Community membership details for difference contact network

### Data Availability

The PDB files for crystal structures used in this study were obtained from RCSB PDB with accession codes 2V0V, 3CQV, 6WMQ, 6WMS, and 8D8I. Details for each PDB accession is provided in Supplementary Table S1. The supplementary movies, data files, and custom codes for plotting and analysis are hosted at https://zenodo.org/doi/10.5281/zenodo.10959054.

## Author Contributions

A.S. conceptualized the project. R.T. and S.S. performed molecular dynamics simulations and molecular docking. V.A.M. and S.S. performed protein contact network analysis. A.S., V.A.M., and S.S. wrote the manuscript.

## Note

The authors declare no conflict of interest.

## Acknowledgements

S.S. acknowledges the support by Council for Scientific and Industrial Research (CSIR), Ministry of Science and Technology, Government of India, for Junior and Senior Research Fellowship. R.T. and V.A.M. acknowledge the funding support from Indian Institute of Technology Gandhinagar, India. A.S. thanks Department of Biotechnology for Ramalingaswami Re-entry Fellowship and Indian Institute of Technology Gandhinagar for internal project grant support.

