## Supplementary Material for "REV-ERBβ Binding Pocket Dynamics with Implications for Rational Design of Small Molecule Modulators"

#### Supplementary Figures

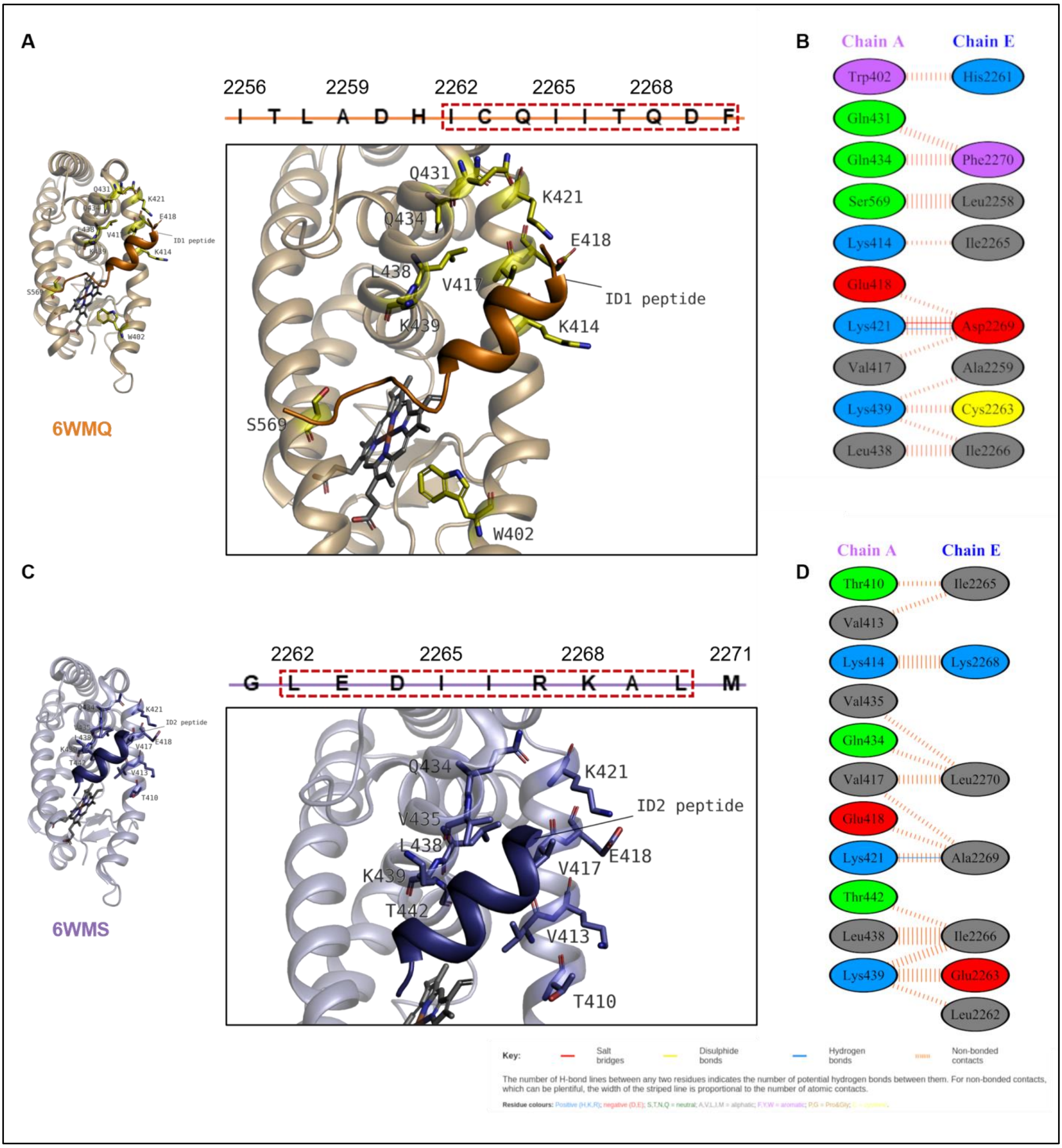


**Supplementary Figure S1: Interaction of ID peptides with REV-ERBβ LBD in presence of heme. (A)** and **(B)** ID1 peptide interactions with REV-ERBβ LBD (PDB ID: 6WMQ). **(C)** and **(D)** ID2 peptide interactions with REV-ERBβ LBD (PDB ID: 6WMS). CoRNR motif is highlighted with red boxes. Additional details about interactions are provided in Table S2B-C. Figures **(A)** and **(C)** rendered in PyMOL. Figure **(B)**, **(D)**, and legend key were taken from PDBsum (Laskowski et al., 2018). Chain A and E refer to REV-ERBβ LBD and ID peptides, respectively.

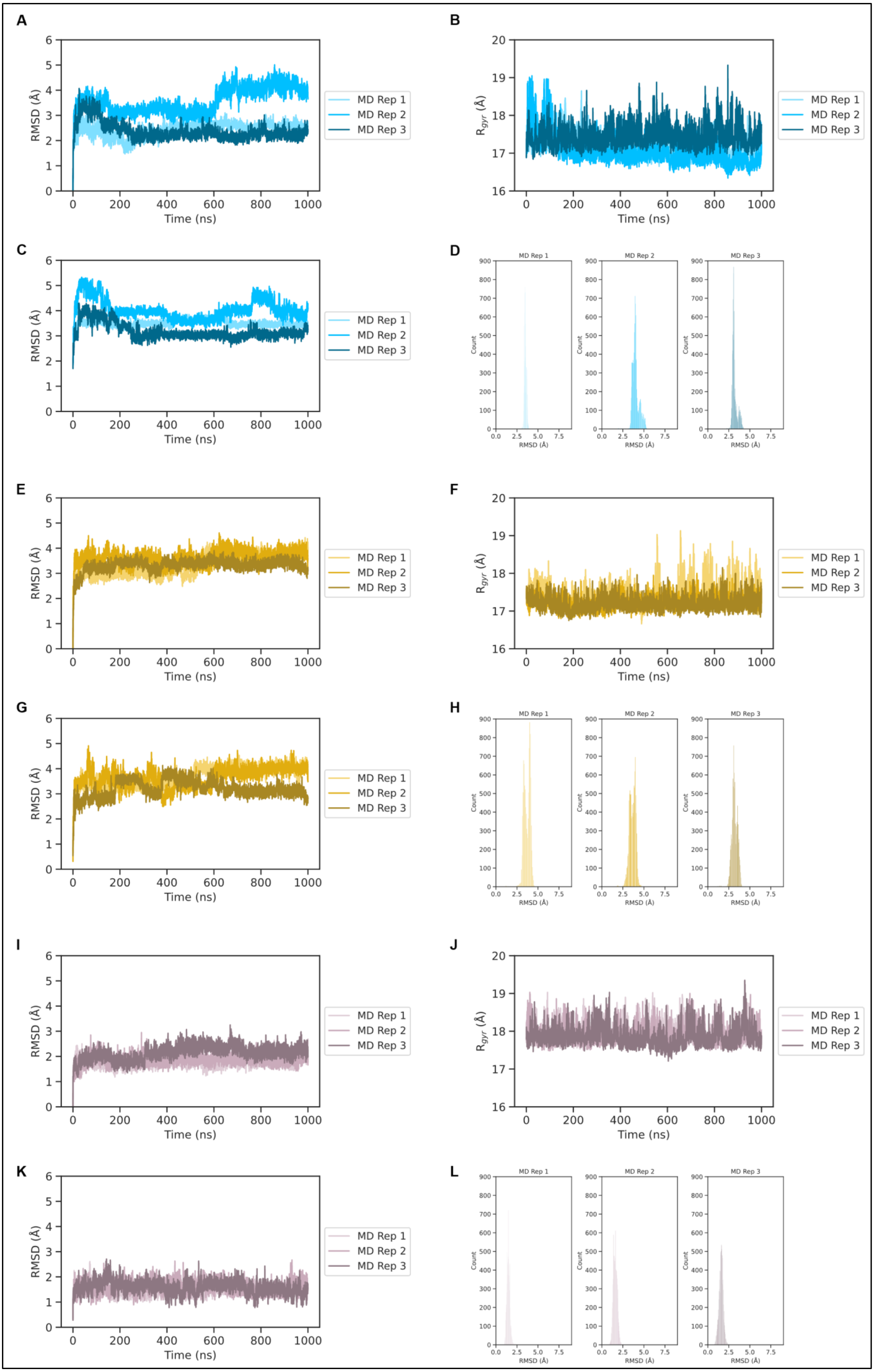

A
RMSD (Å)
Time (ns)
MD Rep 1
MD Rep 2
MD Rep 3
B
$R_{gyr}$ (Å)
Time (ns)
MD Rep 1
MD Rep 2
MD Rep 3
C
RMSD (Å)
Time (ns)
MD Rep 1
MD Rep 2
MD Rep 3
D
MD Rep 1
MD Rep 2
MD Rep 3
Count
RMSD (Å)
E
RMSD (Å)
Time (ns)
MD Rep 1
MD Rep 2
MD Rep 3
F
$R_{gyr}$ (Å)
Time (ns)
MD Rep 1
MD Rep 2
MD Rep 3
G
RMSD (Å)
Time (ns)
MD Rep 1
MD Rep 2
MD Rep 3
H
MD Rep 1
MD Rep 2
MD Rep 3
Count
RMSD (Å)
I
RMSD (Å)
Time (ns)
MD Rep 1
MD Rep 2
MD Rep 3
J
$R_{gyr}$ (Å)
Time (ns)
MD Rep 1
MD Rep 2
MD Rep 3
K
RMSD (Å)
Time (ns)
MD Rep 1
MD Rep 2
MD Rep 3
L
MD Rep 1
MD Rep 2
MD Rep 3
Count
RMSD (Å)

**Supplementary Figure S2: Analysis of 2V0V-A, Apo-3CQVΔN-ter, and Apo-3CQV simulation trajectories (3 replicates). (A) – (D)** Analysis of 2V0V-A trajectories. **(A)** RMSD of REV-ERBβ LBD Cα atoms with $0^{th}$ frame as reference **(B)** Plot for radius of gyration from simulation trajectories. **(C)** RMSD of binding site residue Cα atoms with 3CQV crystal structure as the reference and **(D)** distribution of the RMSD values present in **(C)**. **(E)-(H)** Analysis of Apo-3CQVΔN-ter trajectories. **(E)** RMSD of REV-ERBβ LBD Cα atoms with $0^{th}$ frame as reference **(F)** Plot for radius of gyration from simulation trajectories. **(G)** RMSD of binding site residue Cα atoms with 3CQV crystal structure binding site residue Cα atoms as the reference and **(H)** distribution of the RMSD values present in **(G)**. **(I)-(L)** Analysis of Apo-3CQV trajectories. **(I)** RMSD of REV-ERBβ LBD Cα atoms with $0^{th}$ frame as reference **(J)** Plot for radius of gyration from simulation trajectories. **(K)** RMSD of binding site residues Cα atoms with 3CQV crystal structure binding site residue Cα atoms as the reference and **(L)** distribution of the RMSD values present in **(K).**

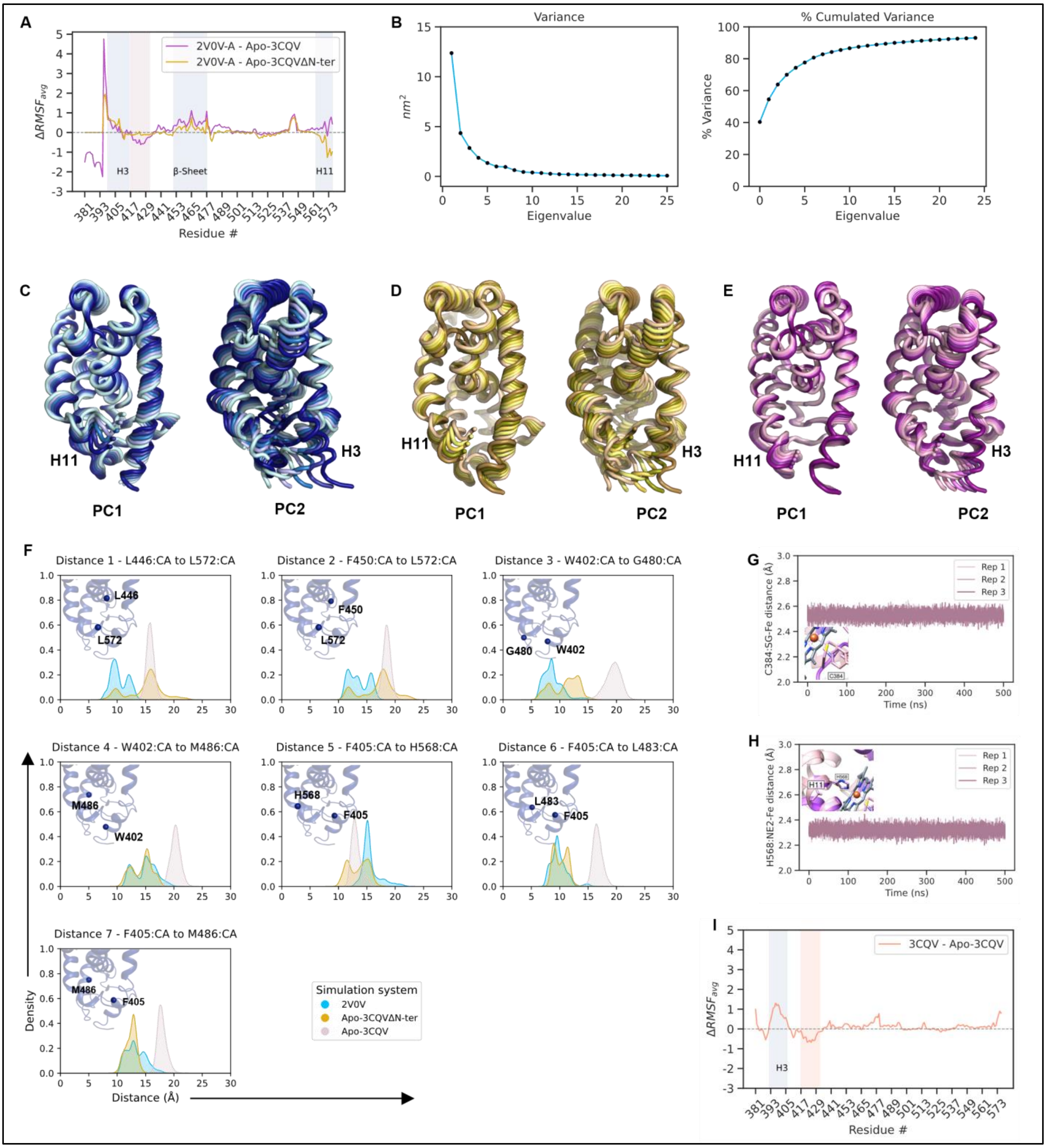


**Supplementary Figure S3: Dynamics of REV-ERBβ LBD in apo and heme bound states. (A)** Difference between average RMSF values (average value from triplicates) in 2V0V-A v/s Apo-3CQVΔN-ter (yellow color), and 2V0V-A v/s Apo-3CQV (magenta color) simulation trajectory. Regions shaded in blue, and pink, represent higher fluctuations in 2V0V-A/Apo-3CQVΔN-ter and Apo-3CQV, respectively. **(B)** Variance explained by top 25 eigenvalues from concatenated simulation trajectory (2V0V-A, Apo-3CQVΔN-ter, and Apo-3CQV, see methods). Points in the graph correspond to the eigenvalues. Global motions in PC1-PC2 space for **(C)** 2V0V-A, **(D)** Apo-3CQVΔN-ter, and **(E)** Apo-3CQV simulation trajectories. The dark to light color gradient represents the motion in the trajectories. Figures **(C)** to **(E)** rendered in PyMOL. **(F)** Distances between Cα atoms of binding site residue pairs in 2V0V-A, Apo-3CQVΔN-ter, and Apo-3CQV simulation trajectories. Distances from 2V0V-A and 3CQV crystal structure are provided in Supplementary Table S3. **(G)** and **(H)** Distance plot between heme iron center ($Fe^{3+}$) and atoms involved in coordinate bonds to stabilize heme binding across 3CQV simulation trajectory. **(I)** Difference between average RMSF values (average value from triplicates) in 3CQV v/s Apo-3CQV simulation trajectory. Regions shaded in blue, and orange represent higher fluctuations in 3CQV and Apo-3CQV, respectively.

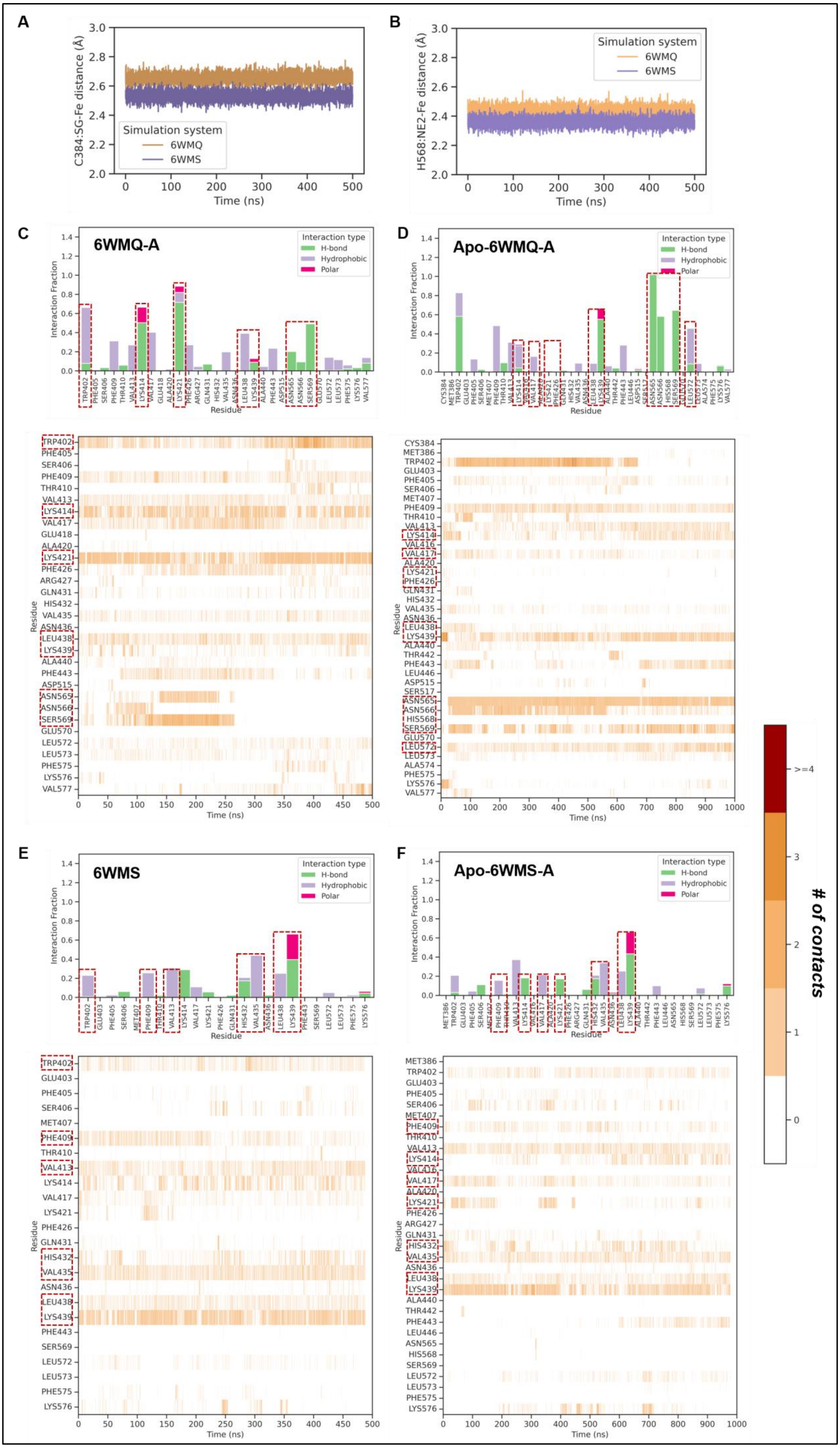

A
C384:SG-Fe distance (Å)
Time (ns)
Simulation system
6WMQ
6WMS
B
H568:NE2-Fe distance (Å)
Time (ns)
Simulation system
6WMQ
6WMS
C
6WMQ-A
D
Apo-6WMQ-A
E
6WMS
F
Apo-6WMS-A
Interaction type
H-bond
Hydrophobic
Polar
Interaction Fraction
Residue
Time (ns)
### of contacts
>=4
3
2
1
0

**Supplementary Figure S4: ID1/2 peptide interactions in presence and absence of heme. (A)** and **(B)** Distance plot between heme iron center ($Fe^{3+}$) and atoms involved in coordinate bonds to stabilize heme binding across ID1 and ID2 peptide bound simulation trajectories (PDB ID: 6WMQ (orange) and 6WMS (purple), respectively). **(C)-(D)** Interactions of ID1 peptide in **(C)** heme bound (6WMQ-A) and **(D)** corresponding apo trajectory (Apo-6WMQ-A). **(E)-(F)** Interactions of ID1 peptide in **(E)** heme bound (6WMS-A) and **(F)** corresponding apo trajectory (Apo-6WMS-A). The bar graph represents the type of interactions and heatmap depicts the contact trend of ID1/2 peptide interactions with REV-ERBβ LBD residues across the simulation trajectories. REV-ERBβ LBD residues forming AF2 surface for ID peptide binding are highlighted in red boxes.

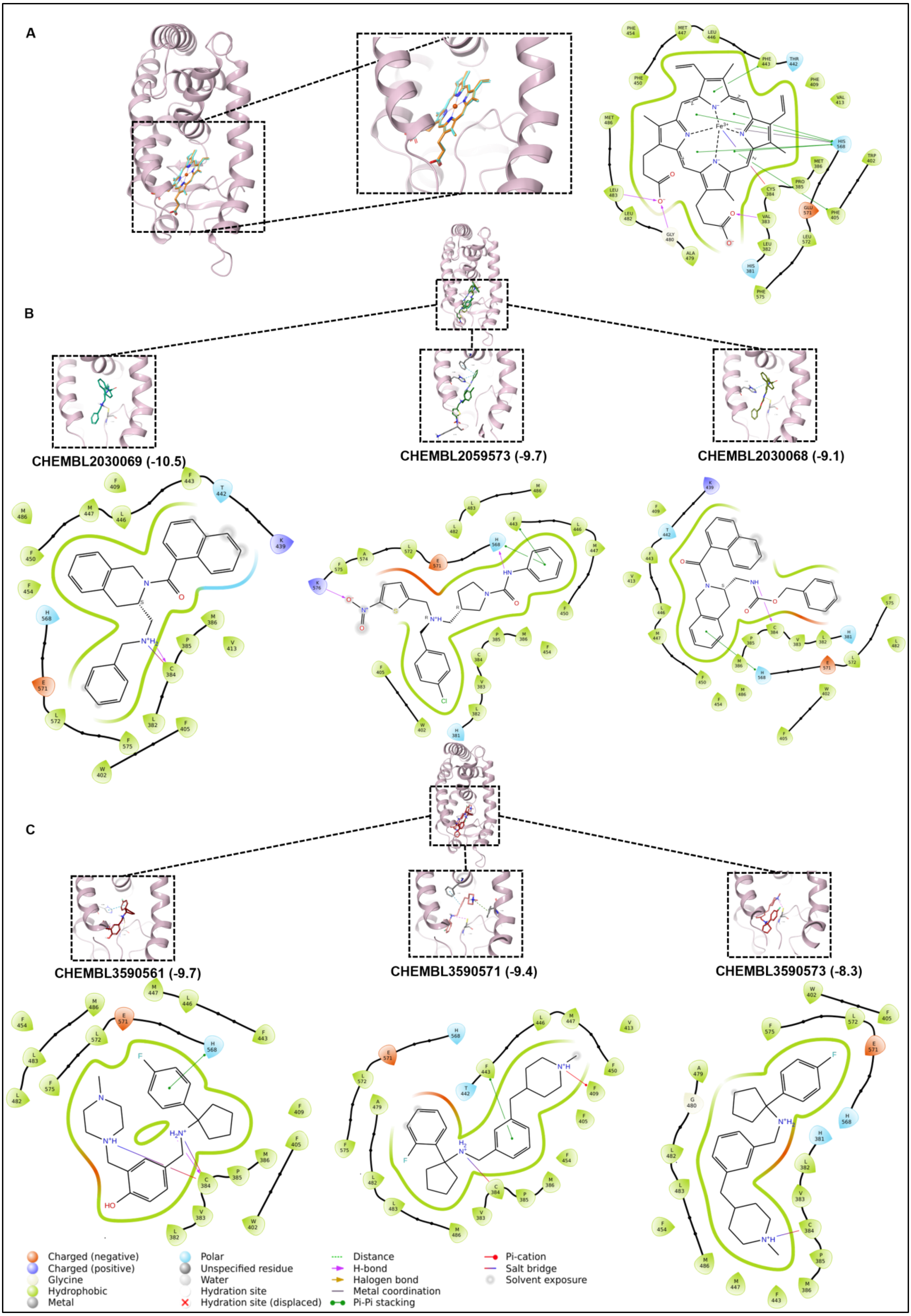

A
B
CHEMBL2030069 (-10.5)
CHEMBL2059573 (-9.7)
CHEMBL2030068 (-9.1)
C
CHEMBL3590561 (-9.7)
CHEMBL3590571 (-9.4)
CHEMBL3590573 (-8.3)
Charged (negative)
Charged (positive)
Glycine
Hydrophobic
Metal
Polar
Unspecified residue
Water
Hydration site
Hydration site (displaced)
Distance
H-bond
Halogen bond
Metal coordination
Pi-Pi stacking
Pi-cation
Salt bridge
Solvent exposure

**Supplementary Figure S5: Binding mode of heme, top three agonists, and antagonists with Apo-3CQV from molecular docking screens. (A)** Binding pose of redocked heme (cyan) to REV-ERBβ (PDB ID: 3CQV) and superposition with the co-crystallized heme (orange) (RMSD: 0.18 Å, docking score: -11.09). Overlay of 3D binding pose of **(B)** top three agonists: CHEMBL2030069, CHEMBL2059573, CHEMBL2030068 and **(C)** top three antagonists: CHEMBL3590561, CHEMBL3590571, CHEMBL3590573. Detailed 3D binding poses and interactions with REV-ERBβ LBD for each small molecule are highlighted in black boxes. 2D ligand interaction diagram highlights the type of interaction with binding site residues (legend key refers to type of residue and interactions). Figures rendered in Schrödinger (Schrödinger Release 2022-3). Docking scores for each small molecule are given in braces. Details of molecular docking of all the small molecules are provided in Supplementary File 1.

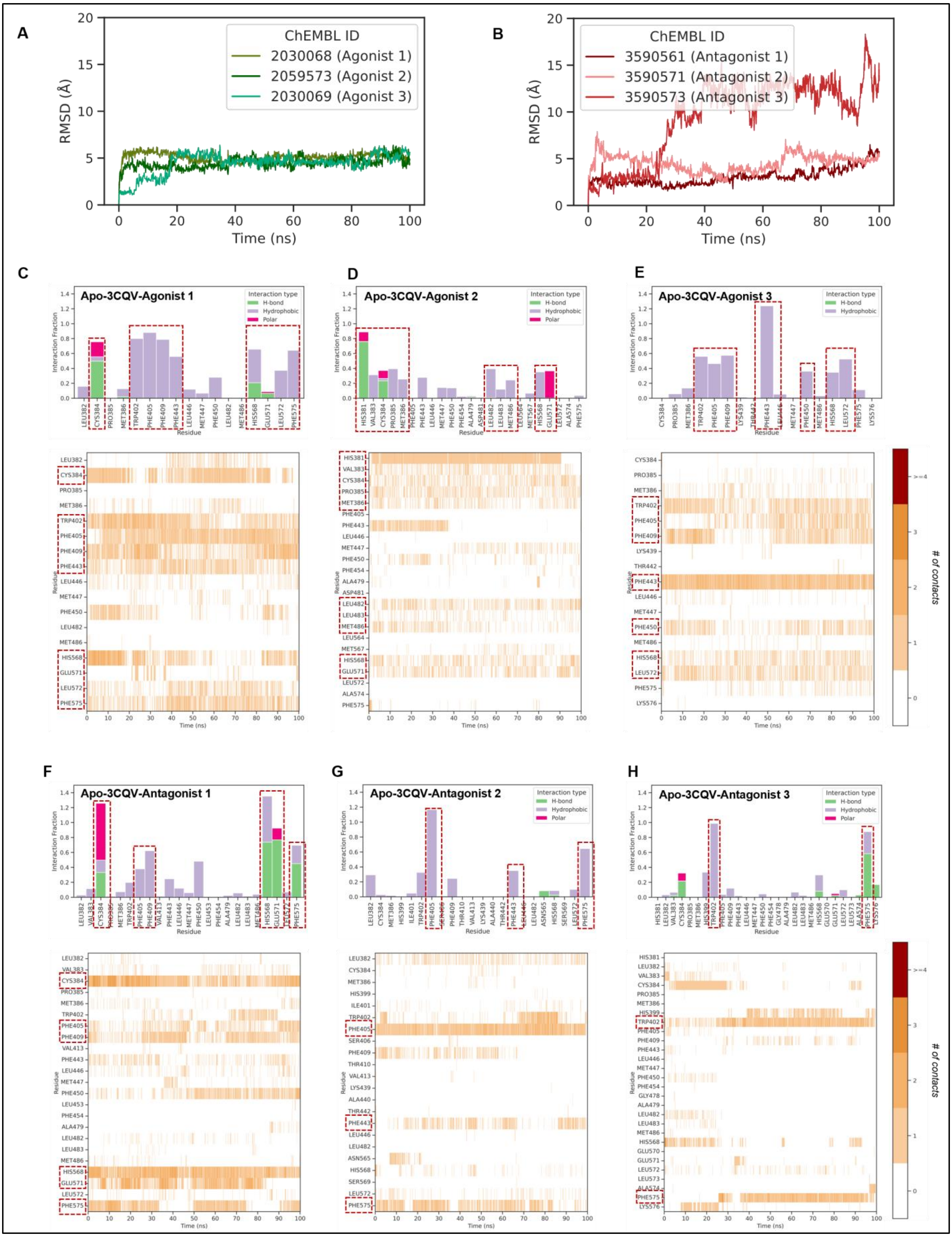


**Supplementary Figure S6: Dynamics of Apo-3CQV-agonist/antagonist docked complexes.** RMSD plot of small molecule **(A)** agonists and **(B)** antagonists in Apo-3CQV simulation trajectories (RMSD calculated w.r.t protein). Interactions of agonists: **(C)** CHEMBL2030069, **(D)** CHEMBL2059573, and **(E)**

CHEMBL2030068 and antagonists: **(F)** CHEMBL3590561, **(G)** CHEMBL3590571, and **(H)** CHEMBL3590573 with REV-ERBβ LBD residues. The bar graph represents the type of interactions and heatmap depicts the contact trend across simulation trajectories. Key binding site residue interactions are highlighted in red boxes.

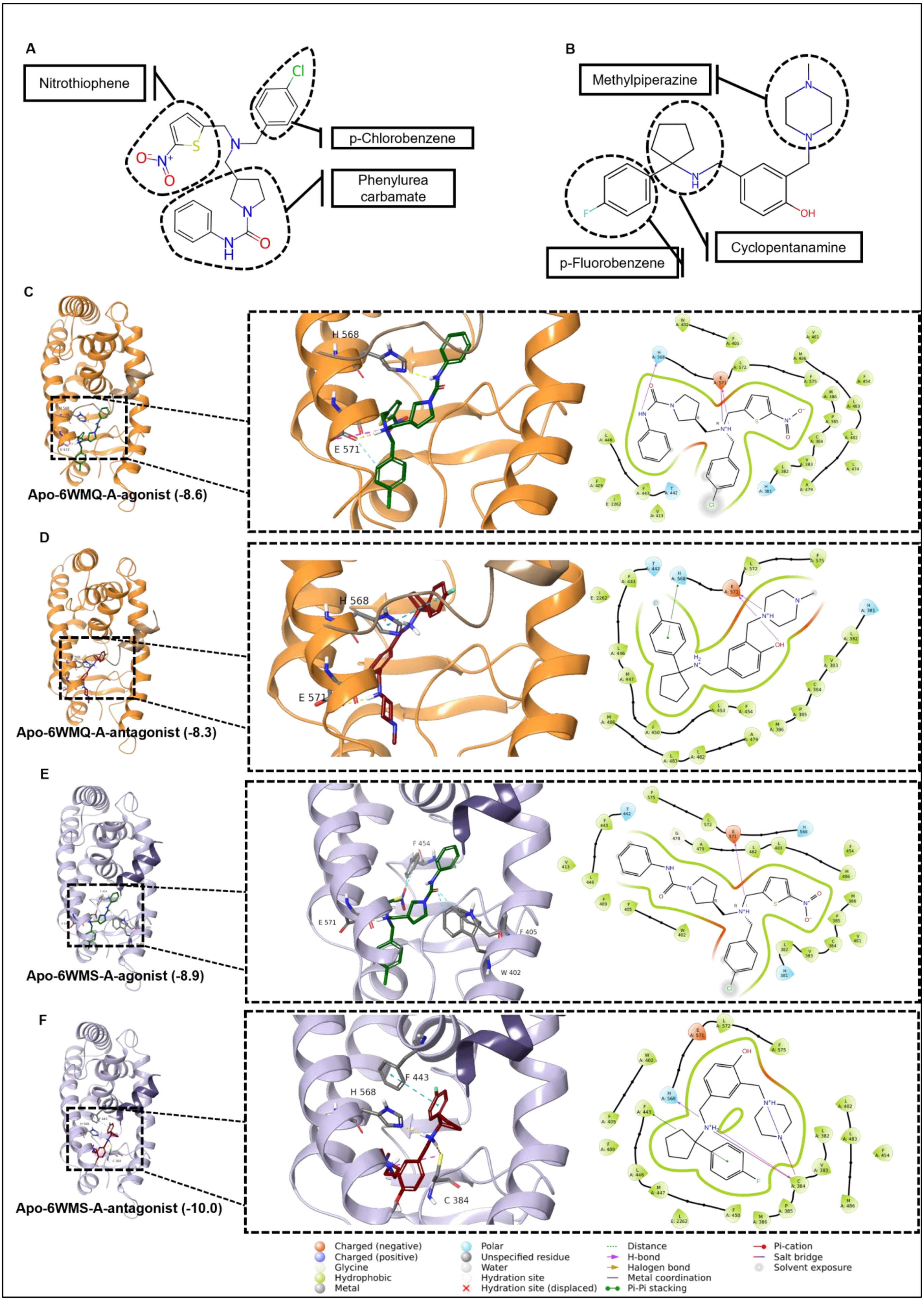

A
Nitrothiophene
p-Chlorobenzene
Phenylurea carbamate
B
Methylpiperazine
Cyclopentanamine
p-Fluorobenzene
C
Apo-6WMQ-A-agonist (-8.6)
H 568
E 571
D
Apo-6WMQ-A-antagonist (-8.3)
H 568
E 571
E
Apo-6WMS-A-agonist (-8.9)
F 454
E 571
F 405
W 402
F
Apo-6WMS-A-antagonist (-10.0)
F 443
H 568
C 384
Charged (negative)
Charged (positive)
Glycine
Hydrophobic
Metal
Polar
Unspecified residue
Water
Hydration site
Hydration site (displaced)
Distance
H-bond
Halogen bond
Metal coordination
Pi-Pi stacking
Pi-cation
Salt bridge
Solvent exposure

**Supplementary Figure S7: Binding modes of agonist and antagonist in ID peptide bound structure.** 2D chemical scaffolds of **(A)** agonist (CHEMBL2059573) and **(B)** antagonist (CHEMBL3590561). Structural features for each small molecule are highlighted. Docked pose of **(C)** agonist (green) and **(D)** antagonist (red) with ID1 peptide bound Apo-6WMQ-A structure (orange). Docked pose of **(E)** agonist (green) and **(F)** antagonist (red) with ID2 peptide bound Apo-6WMS-A structure (purple). 3D binding poses and 2D ligand interactions with REV-ERBβ LBD and ID peptide for each small molecule are highlighted in black boxes (legend key refers to type of residue and interactions). Figures **(C)-(F)** (rendered in Schrödinger (Schrödinger Release 2022-3). Docking scores for each small molecule are given in braces. Details of molecular docking of all the small molecules are provided in Supplementary File 1.

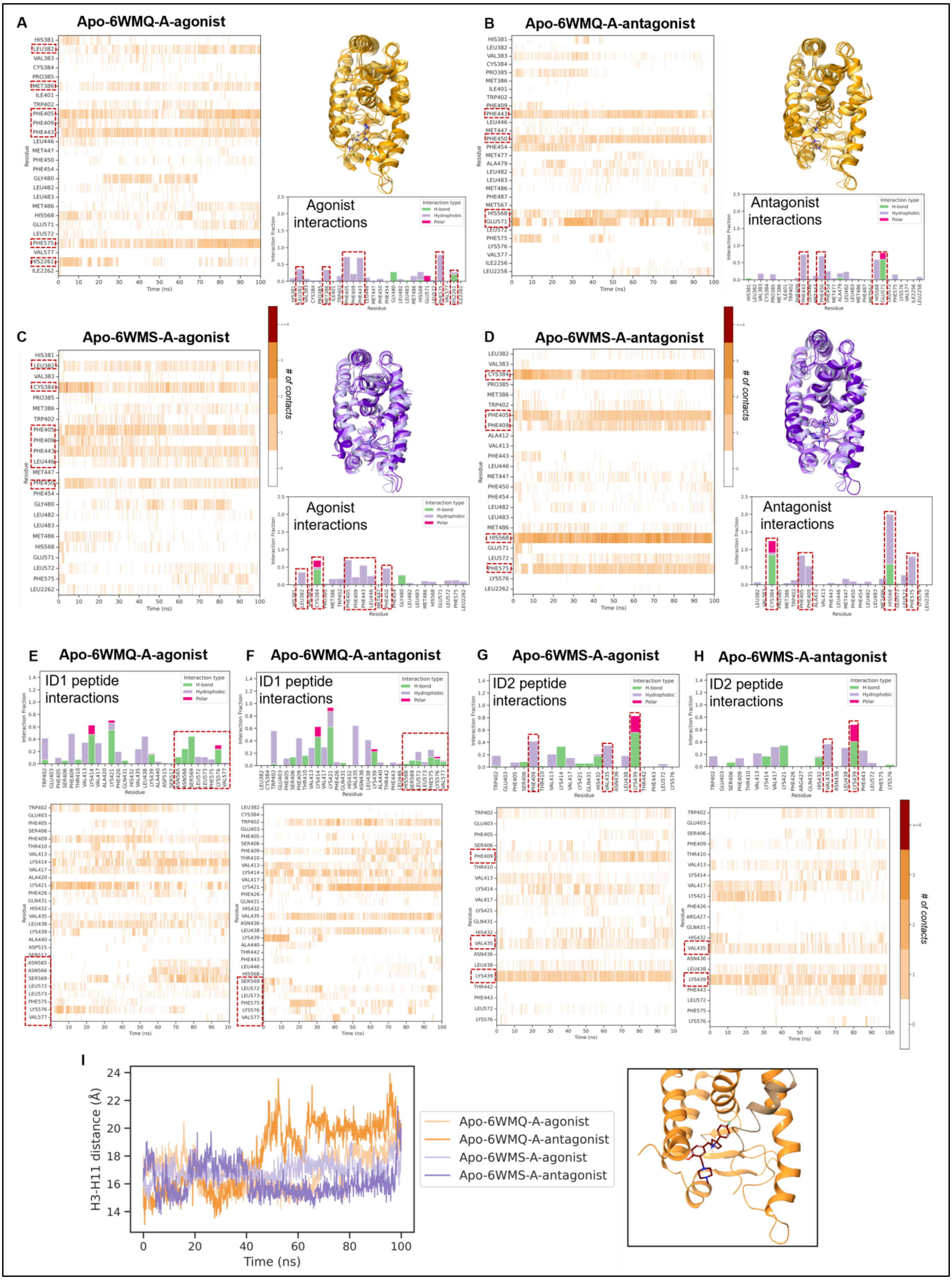


**Supplementary Figure S8: Interactions of agonist (CHEMBL2059573), antagonist (CHEMBL3590561), and ID peptide in simulation trajectories.** Interactions of **(A)** agonist and **(B)**

antagonist with REV-ERBβ LBD in Apo-6WMQ-A-agonist and Apo-6WMQ-A-antagonist simulation trajectories, respectively (ID1 peptide bound). Interactions of **(C)** agonist and **(D)** antagonist with REV-ERBβ LBD in Apo-6WMS-A-agonist and Apo-6WMS-A-antagonist simulation trajectories, respectively (ID2 peptide bound). The trajectory snapshots for each subfigure are colored from light to dark based on the timeline. ID1 peptide interactions with REV-ERBβ LBD residues in presence of **(E)** agonist (Apo-6WMQ-A-agonist) and **(F)** antagonist (Apo-6WMQ-A-antagonist). ID2 peptide interactions with REV-ERBβ LBD residues in presence of **(G)** agonist (Apo-6WMS-A-agonist) and **(H)** antagonist (Apo-6WMQ-A-antagonist). The bar graph represents the type of interactions and heatmap depicts the contact trend across simulation trajectory). **(I)** Distances between Cα atoms of W402-H568 (H3-H11) in Apo-6WMQ-A-agonist (light orange), Apo-6WMQ-A-antagonist (dark orange), Apo-6WMS-A-agonist (light purple) and Apo-6WMS-A-antagonist (dark purple) simulation trajectories. A trajectory snapshot from Apo-6WMQ-A-antagonist trajectory depicting conformational change is highlighted in a black box (rendered in Schrödinger (Schrödinger Release 2022-3)).

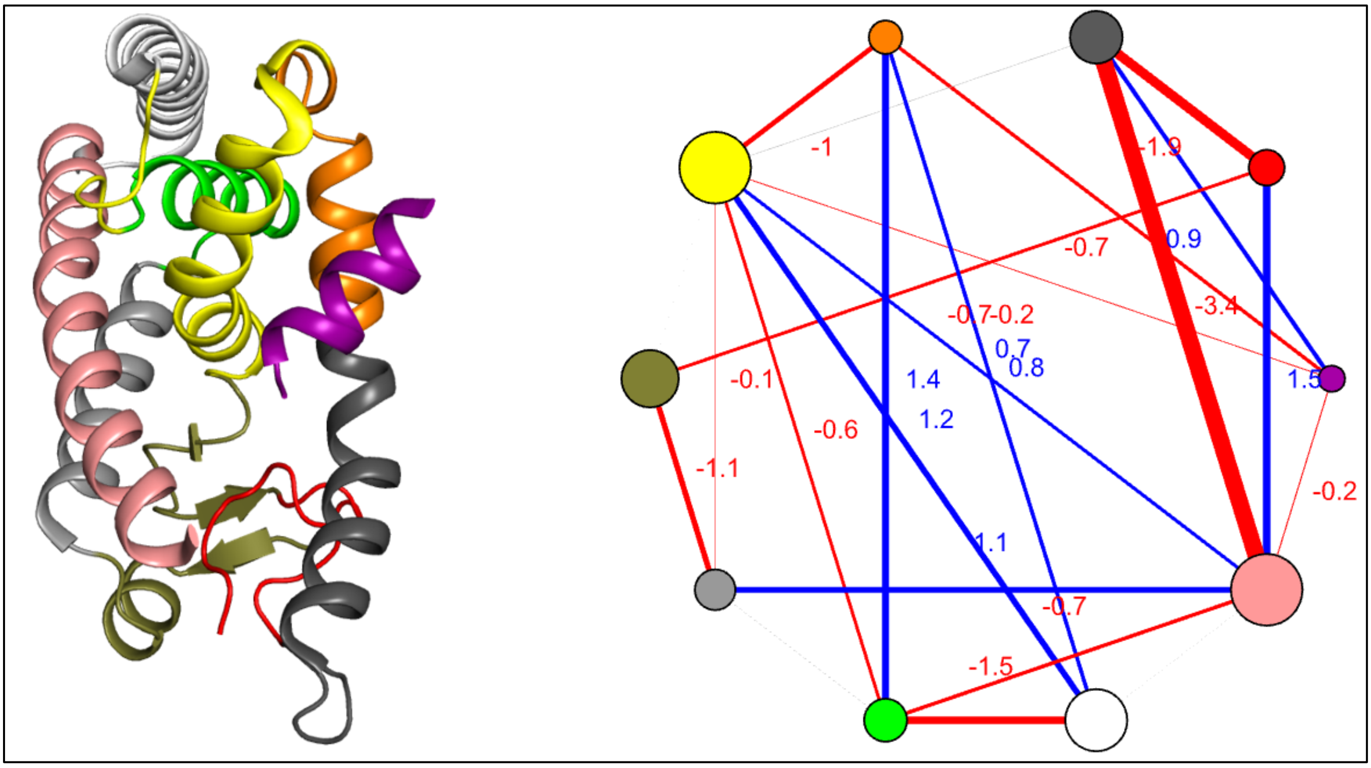


**Supplementary Figure S9:** Apo-6WMS-A-agonist/antagonist (ID2 peptide shown in purple color). Red lines indicate a negative difference, highlighting stronger interactions in the antagonist-bound state, whereas blue lines indicate a positive difference, highlighting stronger interactions in the agonist-bound state. The details of the community membership can be found in Supplementary file 2.

### Supplementary Tables

**Table S1: Crystal structure summary (Uniprot accession: Q14995 (REV-ERBβ) and P20393 (REV-ERBα) )**

| **PDB ID** | **Release date** | **Chains** | **Assembly Composition** | **Resolution (Å)** | **Chain Length** | **Complex** |
|---|---|---|---|---|---|---|
| 6WMQ (Mosure et al., 2021) | 2021-02-17 | A, B, E, F | Heterotetramer | 2.5 | 199 | REV-ERBβ + NCoR ID1 + Heme |
| 6WMS (Mosure et al., 2021) | 2021-02-17 | A, B, E, F | Heterotetramer | 2 | 199 | REV-ERBβ + NCoR ID2 + Heme |
| 4N73 (Matta-Camacho et al., 2014) | 2014-06-04 | A | Monomeric | 1.8 | 199 | REV-ERBβ + Cobalt Porphyrin |
| 3CQV (Pardee et al., 2009) | 2008-08-05 | A | Monomeric | 1.9 | 199 | REV-ERBβ + Heme |
| 2V0V (Woo et al., 2007) | 2007-10-23 | A, B, C, D | Homotetrameric | 2.4 | 194 | REV-ERBβ |
| 2V7C (Woo et al., 2007) | 2007-10-23 | A, B | Dimeric | 2.4 | 194 | REV-ERBβ |

| | | | | | | |
|---|---|---|---|---|---|---|
| 8D8I (Murray et al., 2022) | 2022-12-14 | A | Monomeric | 2.5 | 237 | REV-ERBα + NCoR ID1 + STL1267 |

**Table S2: Interacting residues in REV-ERBβ LBD. (A) Binding Site Residue Correspondence from 3CQV Crystal Structure**

| **Residue** | **Structure** |
|---|---|
| H381, L382, V383, C384, P385, M386 | N-terminal loop |
| W402, F405, F409, V413 | Helix-3 |
| F443, L446, M447, F450 | Helix-5 |
| F454 | β-sheet 1 |
| G478, A479 | Loop before H7 |
| G480, L482, L483, M486 | Helix-7 |
| H568, E571, L572 | Helix-11 |
| F575 | C-terminal loop |

**(B) ID1 Interactions with REV-ERBβ LBD (PDB: 6WMQ) reported in PDBsum** (Laskowski et al., 2018)

| **ID1 Residues** | **LBD Residues** | **LBD Secondary Structure** |
|---|---|---|
| H2261 | W402 | Helix-3 |
| I2265 | K414 | Helix-3 |
| D2269 | V417, E418, K421 | Helix-3 |
| F2270 | Q431, Q434 | Helix-4 |
| I2266 | L438 | Helix-4 |
| A2259, C2263, I2266 | K439 | Helix-4 |
| L2258 | S569 | Helix-11 |

**(C) ID2 Interactions with REV-ERBβ LBD (PDB: 6WMS) reported in PDBsum** (Laskowski et al., 2018)

| **ID2 Residues** | **LBD Residues** | **LBD Secondary Structure** |
|---|---|---|

| | | |
|---|---|---|
| I2265 | T410, V413 | Helix-3 |
| K2268 | K414 | Helix-3 |
| A2269, L2270 | V417 | Helix-3 |
| A2269 | E418 | Helix-3 |
| A2269 | K421 | Helix-3 |
| L2270 | Q434, V435 | Helix-4 |
| I2266 | L438 | Helix-4 |
| I2266 | K439, T442 | Helix-4, Helix-5 |

**Table S3: Distances between Cα atom of binding site residue pair in 2V0V-A and Apo-3CQV crystal structure (in Å). Same distances were measured in 2V0V-A, Apo-3CQVΔN-ter, and Apo-3CQV trajectories.**

| **Residue 1 (Helix #)** | **Residue 2 (Helix #)** | **2V0V-A distance** | **Apo-3CQV distance** | **Δ = Apo-3CQV - 2V0V-A (distance name)** |
|---|---|---|---|---|
| W402 (H3) | M447 (H5) | 15.3 | 20.2 | 4.9 (H3-H5) |
| W402 (H3) | H568 (H11) | 13.2 | 16.6 | 3.4 (H3-H11) |
| L572 (H11) | L446 (H5) | 10.4 | 14.8 | 4.4 (D1) |
| L572 (H11) | F450 (H5) | 13.2 | 17.3 | 4.1 (D2) |
| W402 (H3) | G480 (H7) | 8.7 | 21.5 | 12.8 (D3) |
| W402 (H3) | M486 (H7) | 13.4 | 22.9 | 9.5 (D4) |
| F405 (H3) | H568 (H11) | 13.7 | 16 | 2.3 (D5) |
| F405 (H3) | L483 (H7) | 9.4 | 18.5 | 9.1 (D6) |
| F405 (H3) | M486 (H7) | 11.9 | 20.3 | 8.4 (D7) |

**Table S4: Frequency of Unique Edges in Apo-6WMQ/S-A-ago/antagonist Protein Contact Network**

| Simulation System | Node 1 (chain E) | Node 2 (chain A) | Frequency |
|---|---|---|---|
| Apo-6WMQ-A-agonist | E_2256_ILE | A_440_ALA | 1 |
| | E_2256_ILE | A_515_ASP | 0.02 |
| | E_2256_ILE | A_565_ASN | 1 |
| | E_2256_ILE | A_566_ASN | 1 |
| | E_2257_THR | A_440_ALA | 0.04 |
| | E_2257_THR | A_569_SER | 1 |
| | E_2258_LEU | A_569_SER | 0.8 |
| | E_2258_LEU | A_572_LEU | 0.69 |
| | E_2258_LEU | A_573_LEU | 0.33 |
| | E_2258_LEU | A_577_VAL | 0.06 |
| | E_2260_ASP | A_439_LYS | 0.02 |
| | E_2260_ASP | A_577_VAL | 0.02 |
| | E_2261_HIS | A_577_VAL | 0.12 |
| | E_2265_ILE | A_406_SER | 0.04 |
| | E_2270_PHE | A_434_GLN | 0.2 |

| | | | |
|---|---|---|---|
| Apo-6WMQ-A-antagonist | E_2256_ILE | A_399_HIS | 0.02 |
| | E_2256_ILE | A_402_TRP | 0.04 |
| | E_2256_ILE | A_571_GLU | 0.08 |
| | E_2256_ILE | A_572_LEU | 0.8 |
| | E_2256_ILE | A_573_LEU | 0.12 |
| | E_2256_ILE | A_574_ALA | 0.02 |
| | E_2256_ILE | A_575_PHE | 0.04 |
| | E_2256_ILE | A_577_VAL | 0.04 |
| | E_2257_THR | A_572_LEU | 0.04 |
| | E_2257_THR | A_577_VAL | 0.02 |
| | E_2259_ALA | A_443_PHE | 0.12 |
| | E_2262_ILE | A_413_VAL | 0.75 |
| | E_2265_ILE | A_414_LYS | 0.14 |
| | E_2266_ILE | A_439_LYS | 0.2 |
| Apo-6MWS-A-agonist | E_2261_GLY | A_439_LYS | 0.02 |
| | E_2265_ILE | A_406_SER | 0.96 |

| | | | |
|---|---|---|---|
| | E_2266_ILE | A_410_THR | 0.16 |
| | E_2268_LYS | A_410_THR | 0.02 |
| | E_2269_ALA | A_411_PRO | 0.08 |
| | E_2270_LEU | A_431_GLN | 0.22 |
| | E_2270_LEU | A_435_VAL | 0.24 |
| | E_2271_MET | A_431_GLN | 0.02 |
| Apo-6WMS-A-antagonist | E_2261_GLY | A_576_LYS | 0.24 |
| | E_2263_GLU | A_435_VAL | 0.06 |
| | E_2266_ILE | A_413_VAL | 0.24 |
| | E_2266_ILE | A_414_LYS | 0.02 |

**Table S5: Frequency and Difference Between Common Edges in (A) Apo-6WMQ-A-agonist/antagonist Protein Contact Network**

| Node 1 (chain E) | Node 2 (chain A) | Edge weight (Apo-6WMQ-A-agonist) | Edge weight (Apo-6WMQ-A-antagonist) | Weight difference (agonist-antagonist) |
|---|---|---|---|---|
| E_2256_ILE | A_569_SER | 1 | 0.02 | 0.98 |
| E_2259_ALA | A_439_LYS | 1 | 0.14 | 0.86 |
| E_2259_ALA | A_440_ALA | 0.98 | 0.08 | 0.9 |
| E_2262_ILE | A_439_LYS | 1 | 0.88 | 0.12 |
| E_2263_CYS | A_435_VAL | 0.51 | 0.06 | 0.45 |
| E_2263_CYS | A_439_LYS | 1 | 1 | 0 |
| E_2265_ILE | A_410_THR | 0.43 | 0.04 | 0.39 |
| E_2266_ILE | A_417_VAL | 0.16 | 0.51 | -0.35 |
| E_2266_ILE | A_435_VAL | 0.84 | 0.69 | 0.16 |
| E_2267_THR | A_435_VAL | 0.53 | 0.16 | 0.37 |
| E_2270_PHE | A_431_GLN | 0.73 | 0.31 | 0.41 |
| E_2270_PHE | A_435_VAL | 0.49 | 0.04 | 0.45 |

**(B) Apo-6WMS-A-agonist/antagonist Protein Contact Network**

| Node 1 (chain E) | Node 2 (chain A) | Edge weight (Apo-6WMS-A-agonist) | Edge weight (Apo-6WMS-A-antagonist) | Weight difference (agonist-antagonist) |
|---|---|---|---|---|

| | | | | |
|---|---|---|---|---|
| E_2262_LEU | A_439_LYS | 0.02 | 0.37 | -0.35 |
| E_2263_GLU | A_439_LYS | 1 | 1 | 0 |
| E_2265_ILE | A_410_THR | 0.94 | 0.94 | 0 |
| E_2266_ILE | A_435_VAL | 1 | 0.8 | 0.2 |
| E_2266_ILE | A_439_LYS | 0.18 | 0.75 | -0.57 |
| E_2266_ILE | A_438_LEU | 0.02 | 0.16 | -0.14 |
| E_2267_ARG | A_435_VAL | 1 | 0.76 | 0.24 |
| E_2269_ALA | A_410_THR | 1 | 0.78 | 0.22 |
| E_2269_ALA | A_414_LYS | 1 | 1 | 0 |
| E_2269_ALA | A_413_VAL | 0.2 | 0.53 | -0.33 |
| E_2269_ALA | A_417_VAL | 0.02 | 0.39 | -0.37 |
| E_2270_LEU | A_414_LYS | 0.63 | 0.1 | 0.53 |
| E_2270_LEU | A_417_VAL | 0.1 | 0.96 | -0.86 |

**Table S6: Details of Molecular Dynamics Simulations Performed in Current Study**

| **Simulation System** | **Duration** | **Model Details** |
|---|---|---|
| 2V0V-A | 1μs (3 replicates) | REV-ERBβ LBD apo (chain A) |
| Apo-3CQVΔN-ter | 1μs (3 replicates) | REV-ERBβ LBD apo (residues: 396-576) |
| Apo-3CQV | 1μs (3 replicates) | REV-ERBβ LBD apo |
| 3CQV | 500 ns (3 replicates) | 3CQV crystal structure |
| 6WMQ-A | 500 ns | 6WMQ crystal structure (chain A) |
| 6WMS-A | 500 ns | 6WMS crystal structure (chain A) |
| Apo-6WMQ-A | 1μs | REV-ERBβ LBD apo + ID1 peptide (chain A) |
| Apo-6WMS-A | 1μs | REV-ERBβ LBD apo + ID2 peptide (chain A) |
| Apo-3CQV-CHEMBL2030069 | 100 ns | Apo-3CQV + CHEMBL2030069 |
| Apo-3CQV-CHEMBL2059573 | 100 ns | Apo-3CQV + CHEMBL2059573 |
| Apo-3CQV-CHEMBL2030068 | 100 ns | Apo-3CQV + CHEMBL2030068 |
| Apo-3CQV-CHEMBL3590561 | 100 ns | Apo-3CQV + CHEMBL3590561 |
| Apo-3CQV-CHEMBL3590571 | 100 ns | Apo-3CQV + CHEMBL3590571 |
| Apo-3CQV-CHEMBL3590573 | 100 ns | Apo-3CQV + CHEMBL3590573 |
| Apo-6WMQ-A-agonist | 100 ns | Apo-6WMQ + CHEMBL2059573 |
| Apo-6WMQ-A-antagonist | 100 ns | Apo-6WMQ + CHEMBL3590561 |
| Apo-6WMS-A-agonist | 100 ns | Apo-6WMS + CHEMBL2059573 |
| Apo-6WMS-A-antagonist | 100 ns | Apo-6WMS + CHEMBL3590561 |

**Supplementary Material References**


Laskowski, R. A., Jabłońska, J., Pravda, L., Vařeková, R. S., & Thornton, J. M. (2018). PDBsum: Structural summaries of PDB entries. *Protein Science : A Publication of the Protein Society*, *27*(1), 129–134. https://doi.org/10.1002/pro.3289

Matta-Camacho, E., Banerjee, S., Hughes, T. S., Solt, L. A., Wang, Y., Burris, T. P., & Kojetin, D. J. (2014). Structure of REV-ERBβ Ligand-binding Domain Bound to a Porphyrin Antagonist. *Journal of Biological Chemistry*, *289*(29), 20054–20066. https://doi.org/10.1074/jbc.M113.545111

Mosure, S. A., Strutzenberg, T. S., Shang, J., Munoz-Tello, P., Solt, L. A., Griffin, P. R., & Kojetin, D. J. (2021). Structural basis for heme-dependent NCoR binding to the transcriptional repressor REV-ERBβ. *Science Advances*, *7*(5). https://doi.org/10.1126/sciadv.abc6479

Murray, M. H., Valfort, A. C., Koelblen, T., Ronin, C., Ciesielski, F., Chatterjee, A., Veerakanellore, G. B., Elgendy, B., Walker, J. K., Hegazy, L., & Burris, T. P. (2022). Structural basis of synthetic agonist activation of the nuclear receptor REV-ERB. *Nature Communications*, *13*(1), 7131. https://doi.org/10.1038/s41467-022-34892-4

Pardee, K. I., Xu, X., Reinking, J., Schuetz, A., Dong, A., Liu, S., Zhang, R., Tiefenbach, J., Lajoie, G., Plotnikov, A. N., Botchkarev, A., Krause, H. M., & Edwards, A. (2009). The Structural Basis of Gas-Responsive Transcription by the Human Nuclear Hormone Receptor REV-ERBβ. *PLoS Biology*, *7*(2), e1000043. https://doi.org/10.1371/journal.pbio.1000043

Woo, E.-J., Jeong, D. G., Lim, M.-Y., Jun Kim, S., Kim, K.-J., Yoon, S.-M., Park, B.-C., & Eon Ryu, S. (2007). Structural Insight into the Constitutive Repression Function of the Nuclear Receptor Rev-erbβ. *Journal of Molecular Biology*, *373*(3), 735–744. https://doi.org/10.1016/j.jmb.2007.08.037
